# Regulating Human T Lymphocytes Through Magnetogenetic Tools

**DOI:** 10.1101/2024.10.08.617204

**Authors:** Seyed Hossein Helalat, Rodrigo C. Téllez, Helga Thora Kristinsdóttir, Astrid Dolinger Petersen, Seder Fayyad Islam, Carlos Rodriguez Pardo, Yi Sun

## Abstract

The field of synthetic biology has expanded the possibilities for controlling cellular functions, particularly in the development of mammalian cells for therapeutic applications. This study explored the application of magnetogenetic tools to regulate T cell activity, a crucial aspect of developing advanced immunotherapies. Magnetogenetic tools use magnetic fields to remotely control engineered ion channels and protein domains, providing non-invasive, deep-tissue stimulation that overcomes the limitations of traditional methods. We investigated the effects of three magnetogenetic tools - engineered TRPV1 (TRP1-Fer) and TRPV4 (TRP4-Fer) channels, and Electromagnetic Perceptive Gene (EPG) - in Jurkat cells. First, calcium concentration measurements confirmed the activity of these tools within the cells. Using qPCR and proteomics analysis, we then analyzed their impact on T cell activation, calcium signaling, mitochondrial function, membrane integrity, and gene expression under both stimulated (with antigens) and non-stimulated conditions. Our results revealed significant upregulation of activation and calcium-handling proteins in stimulated cells, indicating enhanced activation and cytoskeletal dynamics compared to controls. However, in non-stimulated cells, the magnetogenetic tools unexpectedly led to deactivation of T cells. This investigation showed that while magnetic induction alone deactivated the cells, antigen stimulation in conjunction with magnetic induction amplified cell activation.

This study highlights the potential of magnetogenetics to precisely modulate T cell functions, presenting promising avenues for more effective and controlled immunotherapies. However, the findings also underscore the need for careful optimization to mitigate potential adverse effects on cellular integrity and function.

## Introduction

Synthetic biology is an interdisciplinary field dedicated to the development of new biological systems and components based on existing knowledge. By applying engineering principles, researchers aim to produce predictable and robust systems with novel functionalities that do not exist in nature. This emerging field bridges biotechnology, biomaterials, and molecular biology, providing essential methodologies and disciplines that advance these areas.^1^

One significant application of synthetic biology is the engineering of mammalian cells for medical purposes. Researchers have created living cell chassis rationally designed from existing signaling networks with new constructs to fulfill various purposes. These include the production of medical biomolecules, the development of synthetic gene networks for sensing or diagnostics, and the programming of organisms to manage disease mechanisms and related biological events.^2^

Cells interact with their environment through a variety of biological signaling systems that impact gene expression. Stimuli perceived by cells can include soluble low-molecular factors such as hormones and growth factors, environmental signals like the extracellular matrix and adhesive molecules, antigens, and physical factors such as mechanical stimuli, temperature changes, or pH alterations. These signals regulate a wide range of cellular activities, including survival, differentiation, migration, and proliferation.^3,4^

The utilization of physical factors as tools for cell function regulation has captivated numerous scientific researchers. Among these physical factors, magnetic fields have gained particular attention due to their strong penetration and non-invasive nature. The technology of magnetic regulation in cells, known as magnetogenetics. This technology typically involves the use of an external magnetic field to stimulate magnetic nanoparticles or ferritin-tagged ion channels. This method impacts cell function and enables the non-invasive regulation of calcium concentrations in cells which is a key signaling messenger.^5^

The quintessential ion channels used in magnetogenetics are transient receptor potential (TRP) vanilloid 1 (TRPV1) or vanilloid 4 (TRPV4) cation channels, which result in the influx of Ca^2+^ when activated. TRPV1 can be activated by heat and ROS, while TRPV4 is activated by mechanical stimuli and also responds to heat. This has led to debates about the effects of regulation and unclear mechanisms of magnetic modulation mediated by these effects.^6,7^

In addition to TRPV1 and TRPV4, EPG (Electromagnetic Perceptive Gene) is another recently emerged magnetogenetics tool that has garnered significant interest. EPG was originally discovered in the glass catfish, *Kryptopterus bicirrhis*, and is highly responsive to electromagnetic fields. When expressed in mammalian cells, EPG can induce significant calcium influx upon exposure to electromagnetic fields, similar to the mechanisms observed in TRP channels. EPG is a membrane-bound protein with its N-terminal domain outside the cell, C-terminal domain inside the cell, and the rest of the protein embedded in or associated with the membrane via a glycosylphosphatidylinositol anchor.^8,9^

One of the key applications of magnetogenetics is in the remote control of neural activity. However, the potential of magnetogenetics extends beyond neural activity control. The external and remote control of signal transduction pathways can provide on-demand and spatial control over cell function, which is crucial in areas such as cancer, cardiovascular disorders, and potentially modern immunotherapy.^10–12^

In modern immunology, T cell engineering has emerged as a groundbreaking field in therapeutic applications, particularly in cancer treatment. The ability to manipulate T cells to recognize and destroy cancer cells has revolutionized immunotherapy. Chimeric Antigen Receptor (CAR) T cell therapy, which involves modifying T cells to express receptors specific to cancer antigens, has shown significant success in treating hematologic malignancies. The engineering of T cells for such therapies involves the integration of synthetic biology techniques to create cells with enhanced efficacy, specificity, and safety profiles.^13^

In recent years, the need to control T cell functions precisely has become increasingly apparent. The tumor microenvironment (TME) presents a highly challenging landscape for T cell function, often characterized by immunosuppressive conditions, low nutrient availability, and physical barriers. As a result, strategies to enhance T cell metabolism, survival, and function within the TME are critical for improving therapeutic outcomes.^14^

Any technology that can upregulate beneficial genes or downregulate detrimental ones provides a highly adaptable approach to optimizing T cell function for various T cell therapies. In these applications, such systems can be programmed to enhance cytotoxicity upon encountering tumor cells or to reduce exhaustion markers, thereby prolonging T cell activity.^15^

The importance of controlling T cell functions extends beyond just enhancing anti-tumor responses. It also includes minimizing potential side effects and improving the overall safety of T cell therapies.^16,17^

In this investigation, we examined TRPV1 and TRPV4 engineered channels as well as EPG magnetogenetic tools in T cells to understand their impact on T cell activity. Our goal was to clarify how magnetic fields can be used to control T cell functions, providing insights into novel therapeutic strategies for diseases where T cells play a crucial role. To achieve this, we expressed and induced these tools in Jurkat cells and analyzed the resulting proteome profile. Additionally, we studied the effects of combining antigen stimulation with magnetic field induction in T cells.

## Results and Discussion

### Magnetogenetics tools expression and assessment

In the context of T cells, magnetogenetics holds promise for a variety of applications. It can be employed to control T cell activation, differentiation, and migration, which are critical processes in both immune responses and immunotherapy. For instance, by remotely controlling signaling pathways within T cells, researchers could potentially enhance the efficacy of T cell-based therapies for cancer or autoimmune diseases.^18^ Additionally, magnetogenetics could be used to fine-tune the behavior of engineered T cells in adoptive cell transfer therapies, improving their specificity and persistence in targeting tumors.^15^ However, the impact of magnetogenetic tools on cellular behavior remains inadequately understood. To address this knowledge gap, we conducted a comprehensive study examining the effects of three different magnetogenetic tools on the proteome profile of T cells. Our research aimed to elucidate how these tools influence T cell function at the molecular level, providing insights into both the potential therapeutic benefits and the underlying mechanisms of magnetogenetic interventions in immune cells. This investigation is a crucial step towards harnessing the full potential of magnetogenetics in synthetic biology and T cell engineering (Figure 1).

**Figure 1.**
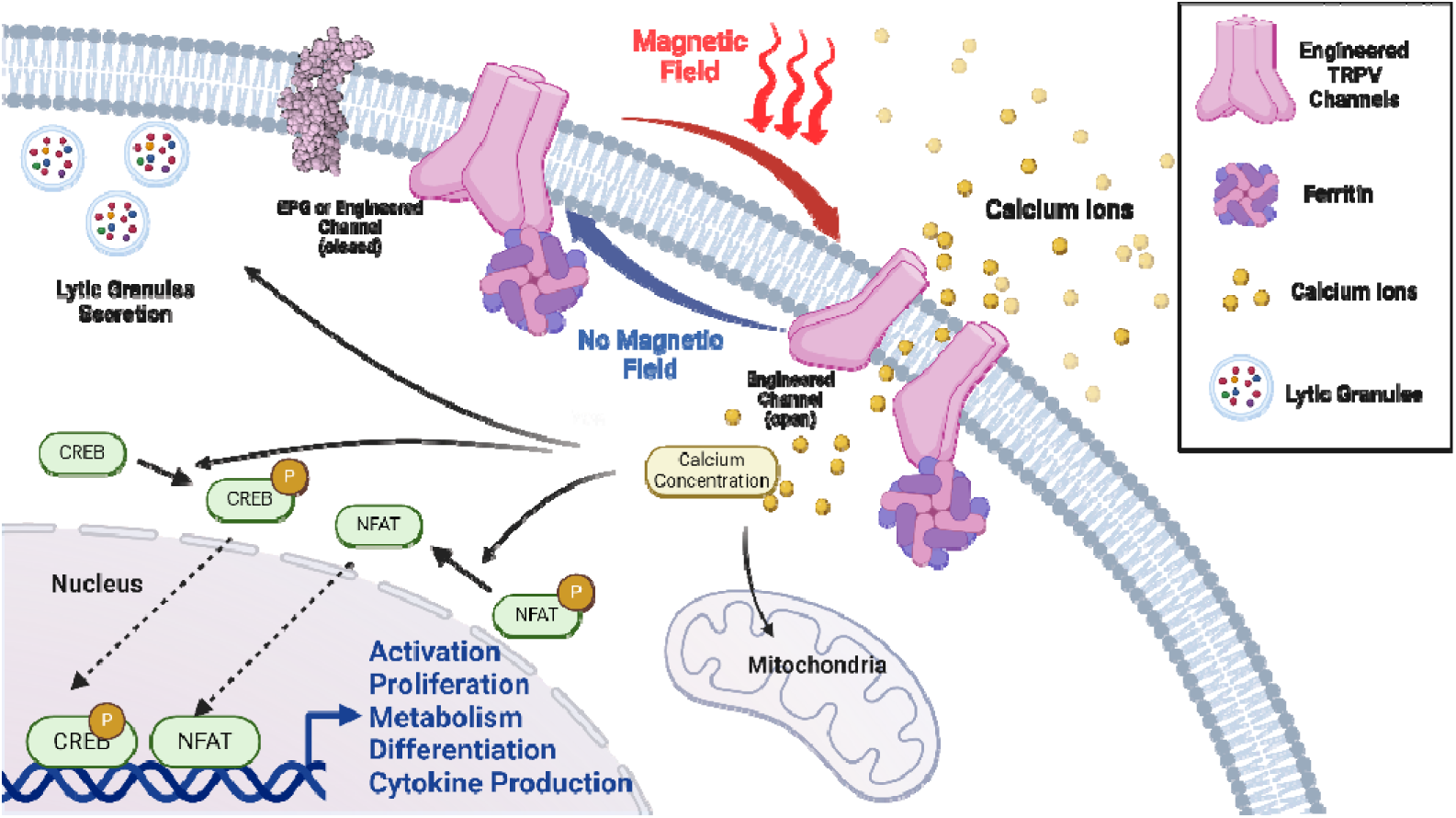
Schematic representation of the effects of magnetogenetic tools on cell behavior. Magnetogenetic tools, upon magnetic field induction, increase intracellular calcium levels by facilitating calcium influx. In T cells, elevated calcium concentration is a crucial regulator of key signaling pathways, particularly those controlling cell activation through the regulation of transcription factors like NFAT. These calcium fluctuations also influence various cellular components, including mitochondria, the cell membrane, and ion-regulating proteins, leading to broader physiological effects.

In this study, the chosen tools were TRPV1 and TRPV4 engineered channels as well as EPG protein domain. The utilized engineered TRPV1 (TRP1-Fer) was a TRPV1 channel fused with an anti-GFP nanobody, which interacts with engineered ferritin containing an integrated GFP-tagged ferritin dimer subunit.^6^ This complex, when exposed to a magnetic field, leads to Ca^2+^ influx, which can control Ca^2+^-dependent cell signaling. The other TRPV-derivated tool was Magneto2.0 (TRP4-Fer). This tool was engineered by fusing the cation channel TRPV4 to the paramagnetic protein ferritin, allowing the channel to be activated by magnetic fields.^7^

Moreover, we chose to use the EPG protein domain, which offers a novel method for controlling cellular functions using a magnetic field. This protein does not require any additional fused ferritin or magnetically inducible nanoparticles and has a short sequence, making it a promising tool for T cell engineering via lentiviruses, which have limitations in gene size.^8^

To examine the tools being induced by magnetic field. First, we made Minicircle plasmids harboring EPG, TRP1-Fer, and TRP4-Fer genes to confirm the functionality of the created tools, we transfected Hek cells. Afterward, the cells were induced by a moving permanent magnet, and the concentration of calcium was measured. The calcium concentration assay revealed the functionality of the engineered TRPV channels and EPG in increasing calcium concentration in Hek cells (Figure 2A). Moreover, fluorescence imaging was utilized for real-time observation of the calcium concentrations increased by the system (Figure 2B).

**Figure 2.**
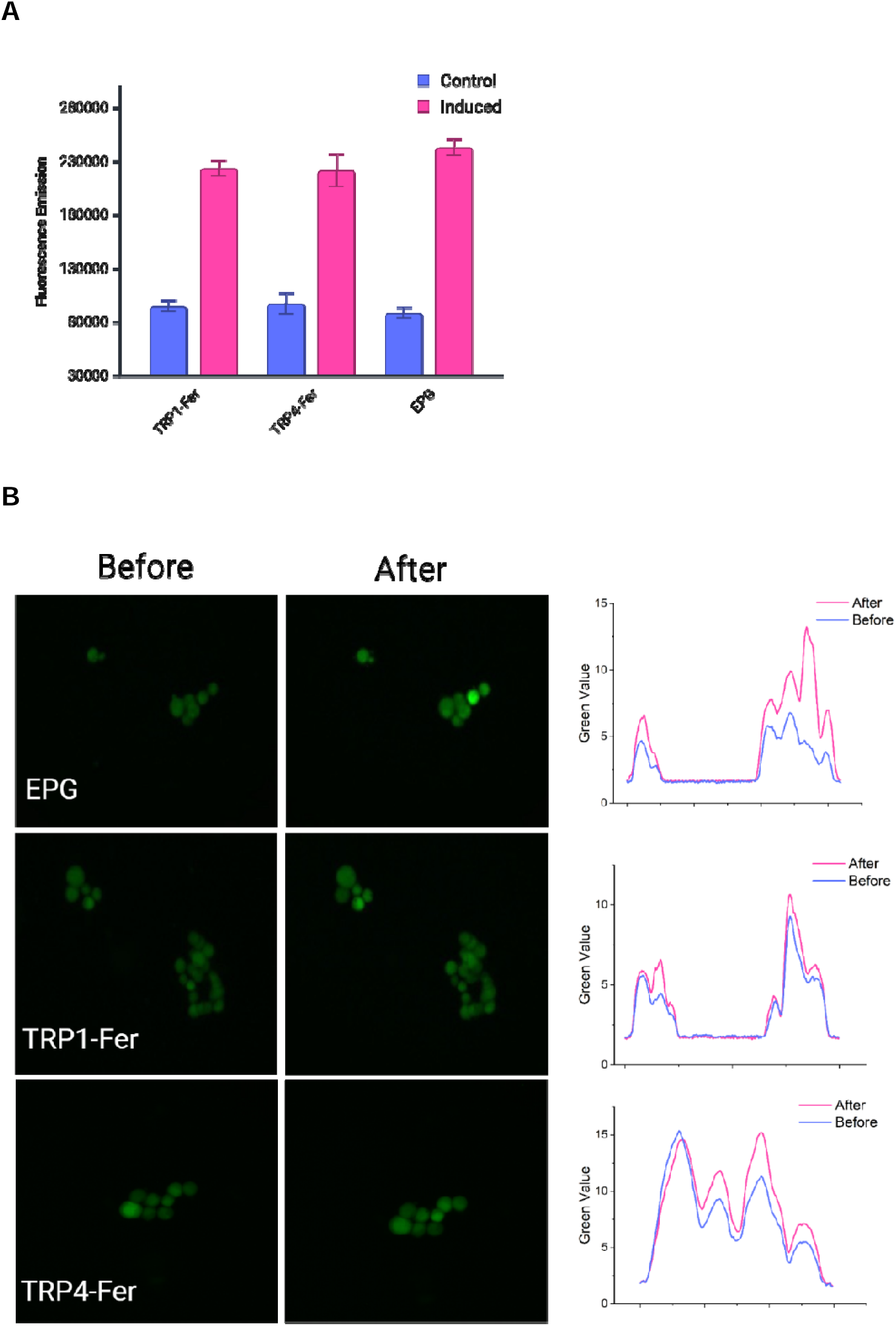
Calcium concentrations measurement of induced tools expressing cells and viability assay of tools expressing cells. (A) Fluorescent calcium indicators demonstrated an increase in calcium levels in Hek cells expressing various tools after induction. (B) Fluorescence microscopy images of Hek cells, with different tools, were captured while using a moving permanent magnet to observe calcium concentrations post-induction. The green fluorescence intensity was quantified using ImageJ software to better illustrate signal fluctuations.

Then, we generated lentiviruses harboring engineered EPG, TRP1-Fer, and TRP4-Fer genes to create stable cell lines, thereby eliminating the effects of episomal expression in the experiments. These constructs were designed to express the mCherry gene for EPG and TRP4-Fer, and GFP for TRP1-Fer. The lentiviruses were used to transduce Jurkat cells, a well-known model for T cell studies, followed by sorting the gene-expressing positive cells using FACS. The toxicity of the construct expression was assessed over 7 days of continuous culture of the sorted positive cells. This experiment showed no impact on cell viability or proliferation compared to normal cells (Figure 3A).

**Figure 3.**
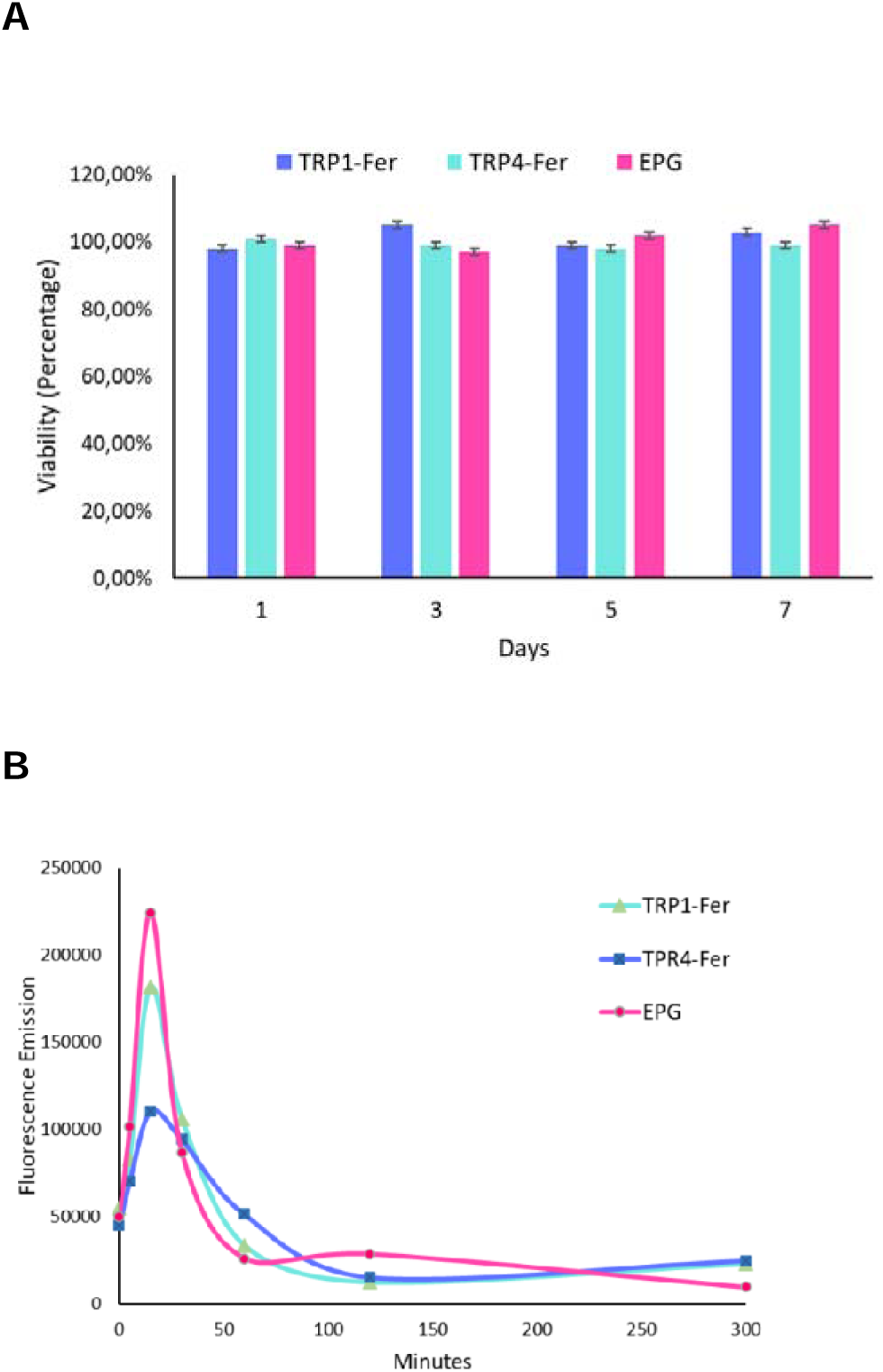
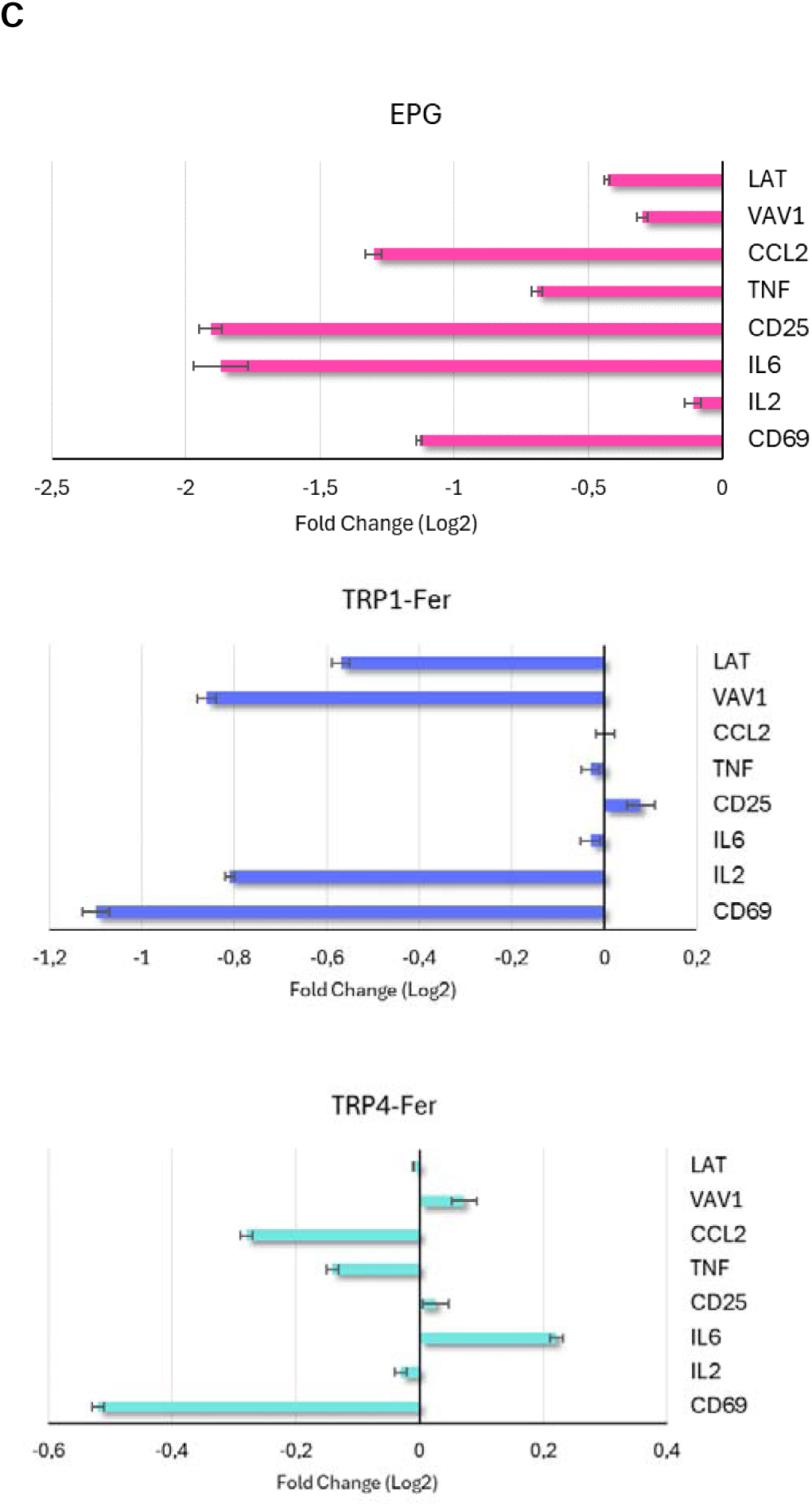
The effect of the tools on Jurkat cells. (A) Viability of Jurkat cells expressing the tools over 7 days. No toxicity was observed from tool expression. (B) Calcium concentration measurement in Jurkat cells expressing the tools over 5 hours of magnetic induction. (C) qPCR results of the tools-expressing cells after 6 hours of magnetic induction. Indicating that induction of the tools could deactivate Jurkat cells.

The sizes of the constructs for virus production varied due to the different tool sizes. The EPG+mCherry construct had a shorter segment of 1200 bp, resulting in higher virus production efficiency compared to the TRPV channels, which had sizes of 4887 bp and 4440 bp for TRP1-Fer and TRP4-Fer, respectively (Table S1). Additionally, we assessed the stability of gene expression in transduced cells for these constructs after 120 days of culture. The EPG construct showed 95% stability of expression, while TRP4-Fer and TRP1-Fer had 84% and 72% positive expressing populations, respectively. This variation in stability could be due to the larger sizes of these genes.

In this investigation, we tested the use of an alternating magnetic field (AMF) generated by electromagnets, alongside a moving magnetic field, to induce T cells. Although AMF induction had a stronger effect on the cells, it increased the culture temperature by 10-12 degrees Celsius (Figure S1A). Consequently, we focused more on using moving permanent magnets (MPM) due to the thermal effects of AMF. We established and evaluated two methods: one involved a fixed cell plate (FCP) with a moving row of magnets beneath it, and the other involved shaking the cells on a fixed magnet plate (FMP), which could only be used for suspension cells (Figure S1B and C). No significant difference was observed between using FCP and FMP in our setup, and they were interchangeable based on convenience.

### Calcium concentration and qPCR assessments

To study the effects of these tools on Jurkat cells, we first analyzed the changes in calcium concentrations over a period of 6 hours. Our results revealed that calcium levels initially increased within the first 15 minutes but then dropped significantly. This fluctuation suggests a negative feedback mechanism in response to the initial rise in calcium concentrations induced by the system (Figure 3B).

To further investigate this phenomenon, we examined the expression levels of key genes in T cell activation including CD69, an early activation marker in T cells, as well as IL-2 and IL-6,^19^ within the same 6-hour induction period. These analyses were performed to assess the impact of the tools on cell function at the transcriptomic level. Our results demonstrated that magnetic induction led to a decrease in the expression of CD69, IL-2, and IL-6 in the induced cells. Additionally, we evaluated the expression of other key genes, including CD25, TNF, CCL2, VAV1, and LAT,^20,21^ all of which indicated that the induction system is steering the cells towards a deactivated state (Figure 3C).

Then, we further investigated the effects of our system in conjunction with low concentrations of anti-CD3 and anti-CD28 stimulatory antigens. To this end, we added antibodies to the culture of the TRP1-Fer, TRP4-Fer, EPG, and control, then induced the cells with the magnetic field. The findings indicated that the expression of CD69, IL2, and IL6, increased compared to the control cells, which were exposed to a magnetic field and antigens (without tool expression). Next, we explored additional genes involved in calcium signaling and regulatory pathways in T cells to gain a deeper understanding of the system’s effects (Figure 4A).

**Figure 4.**
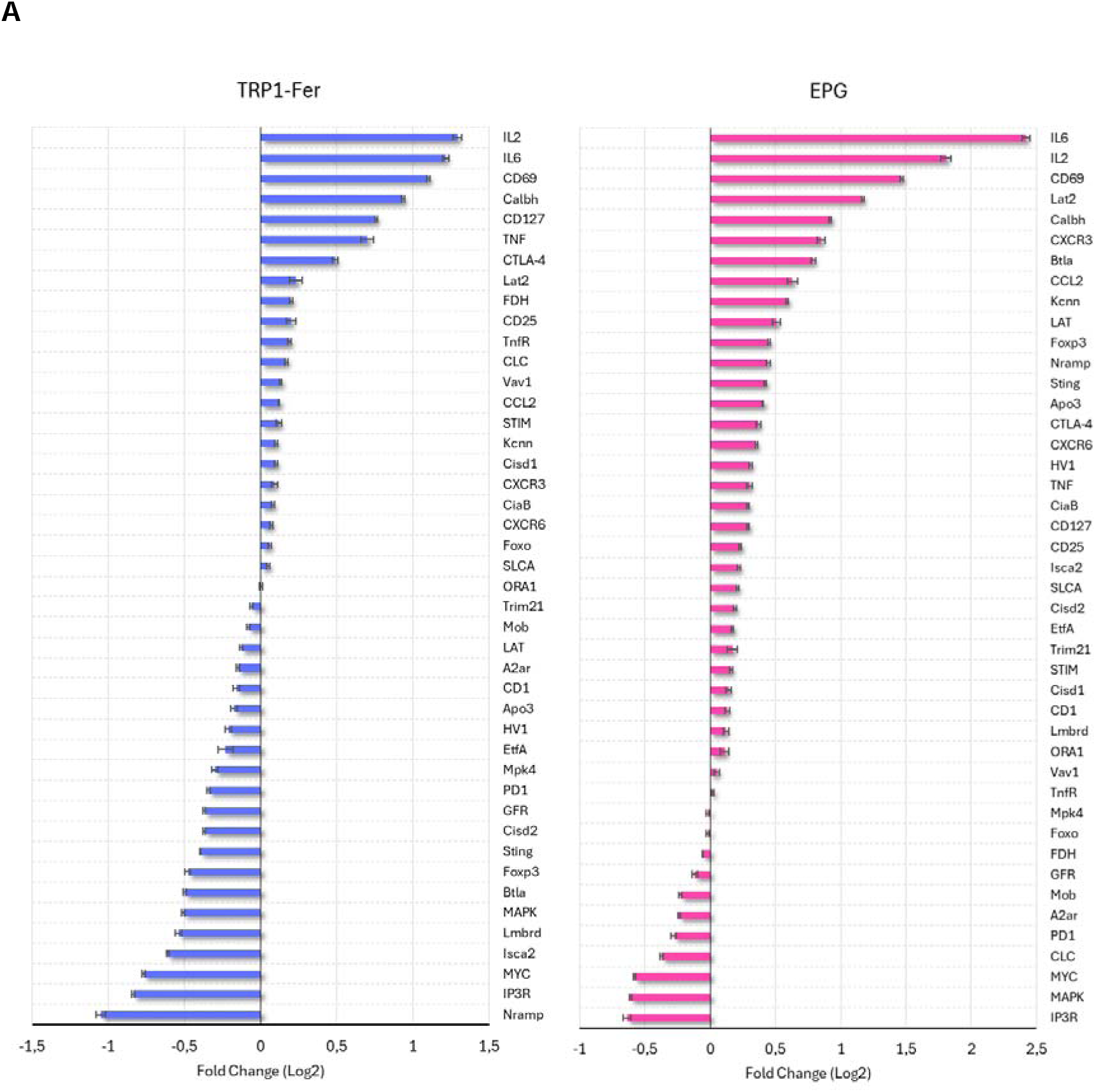
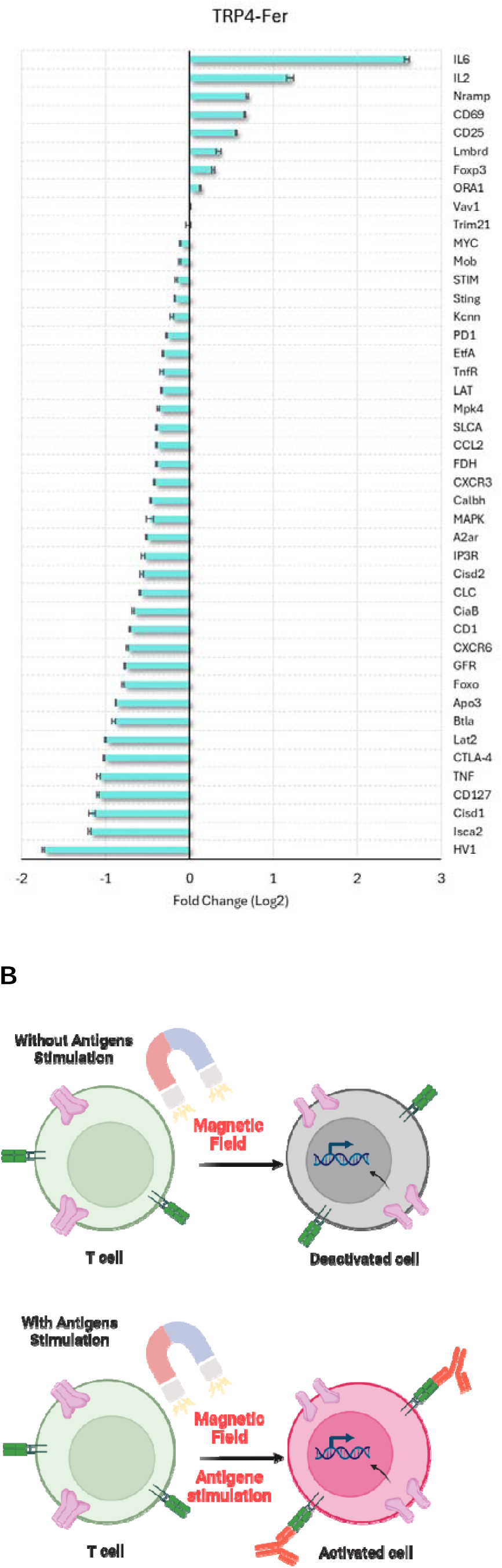
Effect of tools induction on cells coupled with antigens stimuli. (A) qPCR results of Jurkat cells expressing the tools after 6 hours of magnetic induction and stimulation with anti-CD3 and anti-CD28 antibodies, compared to normal control cells without expressing the tools. (B) With the employed tools and the induction setup, the induction led to the deactivation of Jurkat cells. However, when anti-CD3 and anti-CD28 antibodies were added, the magnetic induction amplified the activation of the cells.

The expression of CD69, CD25, IL2, and IL6—markers of T cell activation^19,21^—showed a significant increase in cells exposed to magnetogenetic tools compared to the control group. CD69 is an early activation marker, and its elevated levels indicated that the magnetogenetic induction coupled with antigen stimulation successfully enhanced T cell activation. CD25, another key marker of T cell activation, was also upregulated. IL2 and IL6 are crucial cytokines in T cell proliferation and differentiation, and their increased expression suggests that magnetogenetic tools may enhance the immune response by promoting cytokine production. CD127 (IL-7 receptor alpha) was upregulated in TRP1-Fer and a bit in EPG, suggesting enhanced responsiveness to IL-7 signaling, which plays a crucial role in T cell survival and homeostasis. However, a decrease was observed in TRP4-Fer.

CTLA-4 and PD1 are inhibitory receptors that regulate T cell activation to prevent overactivation and autoimmunity.^22^ While CTLA-4 was upregulated by TRP1-Fer and EPG, PD1 was downregulated in all cells, for promoting sustained T cell activity. This balance between activation (increased CD69, IL2, IL6) and inhibition (modulation of CTLA-4 and PD1) highlights the fine-tuning of T cell responses under magnetogenetic influence. Albeit CTLA-4 expression in TRP4-Fer was decreased. BTLA, another inhibitory receptor that promotes immune tolerance,^23^ showed a more variable response across the tools. The downregulation of BTLA in TRP1-Fer and TRP4-Fer points to a decreased inhibitory tone, which could enhance T cell activation by reducing suppression. This could lead to a more robust immune response in these cells. In contrast, BTLA was upregulated in EPG-expressing cells, suggesting a compensatory increase in inhibitory signaling to counterbalance heightened cell activation. This upregulation likely serves to prevent excessive immune activation and promote immune homeostasis. The differences in BTLA regulation between tools emphasize that the magnetogenetic tools exert distinct effects on T cell signaling pathways, modulating the balance between activation and inhibition to optimize immune responses for specific conditions.

In the examination of the calcium signaling pathway, several key genes involved in calcium release and store-operated calcium entry (SOCE) showed differential expression patterns. IP3R was downregulated, potentially indicating a feedback mechanism to limit excessive calcium release from intracellular stores,^24^ which could be a protective response to maintain cellular calcium homeostasis and prevent calcium overload. ORA1 (also known as Orai1) and STIM, both crucial components of SOCE, remained stable in their expression levels. This stability suggests that under the conditions tested, the regulation of calcium influx remains tightly controlled, ensuring that sustained calcium signaling, essential for various cell functions, is maintained without triggering aberrant calcium responses.^25^ Moreover, the slight upregulation and downregulation of KCNN, a calcium-activated potassium channel,^26^ indicates an adaptive mechanism to help stabilize the membrane potential during prolonged periods of calcium influx. This regulation likely supports sustained cell activation by preventing depolarization-induced calcium entry, thus maintaining the T cells in an active state capable of responding effectively to antigenic stimulation. Moreover, Calcineurin B Homologous Protein 3 (Calbh), a key regulator in calcium-dependent signaling pathways,^27^ was upregulated in both TRP1-Fer and EPG-expressing cells but downregulated in TRP4-Fer. The increased expression of Calbh in TRP1-Fer and EPG indicates heightened calcium-mediated signaling, which could strengthen cell activation, enhance proliferation, and sustain the immune response.

In addition, several key ion-related genes, including CLCN3 (CLC), NRAMP, Voltage-gated hydrogen channel 1 (Hv1), and SLC4A (Solute carrier family 4), showed differential expression patterns across the three magnetogenetic tools, reflecting distinct impacts on ion transport and cell function. CLC, involved in chloride ion (H^+^/Cl^-^) transport,^28^ exhibited almost stable expression in TRP1-Fer cells but was downregulated in both TRP4-Fer and EPG-expressing cells. Its downregulation in TRP4-Fer and EPG cells could indicate a reduced capacity for chloride homeostasis, potentially affecting T cell signaling and function by altering ion flux and cellular excitability. NRAMP, crucial for metal (Fe^2+^, Mn^2+^, etc.) transport and homeostasis,^29^ displayed contrasting expression patterns: it was downregulated in TRP1-Fer but upregulated in both TRP4-Fer and EPG-expressing cells. Hv1, essential for regulating intracellular pH during immune activation,^30,31^ was downregulated in both TRP1-Fer and TRP4-Fer but slightly upregulated in EPG-expressing cells. it is reasonable to hypothesize that the differential expression of Hv1 is a response to changes in intracellular pH modulated by magnetogenetic tools. This could be a protective mechanism to maintain cellular homeostasis, enabling cells to function effectively under the metabolic demands and stress conditions introduced by magnetic field stimulation. Also, SLCA, a bicarbonate transporter involved in pH regulation,^32^ remained almost stable in both TRP1-Fer and EPG-expressing cells, with a slight downregulation in TRP4-Fer.

The expression of C-C motif chemokine 2 (CCL2) and receptors such as CXCR3 and CXCR6, both critical for T cell migration and trafficking,^33–35^ showed differential regulation patterns across the different tools. These findings suggest that the effects of magnetogenetic tools on the migratory potential of T cells vary, potentially influencing how T cells respond to inflammatory signals and migrate toward target tissues.

This observation implies that these tools could modulate T cell trafficking and tissue-specific homing in a context-dependent manner, potentially impacting immune surveillance and response to inflammation or infection. Also, APOBEC-3C (Apo3), a gene involved in viral defense and cytidine deamination,^36,37^ showed a similar pattern of regulation, indicating that the magnetogenetic tools might also affect T cell antiviral responses.

Furthermore, the MAPK signaling pathway plays a critical role in T cell activation and immune response regulation. For MAPK1 (Mitogen-Activated Protein Kinase 1), we observed consistent downregulation across all tools. This showed that the magnetogenetic tools effect could suppress MAPK1 activity, potentially dampening T cell receptor signaling and subsequent effector functions. MAPK1 is involved in the downstream signaling cascade that leads to cytokine production and cell survival; thus, its reduced expression could modulate the extent of T cell activation.^38,39^ Besides, MPK4 (Mitogen-Activated Protein Kinase 4) is part of the stress response pathway and influences cytokine production and T cell differentiation.^40^ MPK4 was slightly downregulated across all tools. Although this change is mild, it could still indicate a subtle shift in the cellular stress response and regulation of inflammatory processes. A decrease in MPK4 might suggest a reduction in the MAPK-mediated anti-inflammatory signaling, potentially affecting the overall balance of immune activation and resolution.

MYC, a crucial regulator that promotes cell proliferation and survival in response to TCR (T cell receptor) and IL-2 signaling,^41^ was consistently downregulated. This reduction in MYC expression suggests a shift towards a less proliferative or more differentiated T cell state under magnetogenetic stimulation.

Additionally, LAT (Linker for Activation of T cells) and LAT2 (Linker for Activation of T cells family member 2), key adaptor molecules essential for TCR (T cell receptor) signaling and downstream calcium mobilization were studied.^42,43^ Our results exhibited a dynamic expression pattern across the different magnetogenetic tools in these adaptor proteins. Foxp3, the master regulator of regulatory T cells (Tregs) and a factor to suppress the function of NFAT and NFkappaB,^44^ showed a mixed response. It was downregulated in TRP1-Fer, stable in TRP4-Fer, and upregulated in EPG. On top of that, FoxO1 (Forkhead Box O1) is a transcription factor that regulates T cell homeostasis and memory formation. FoxO1 expression remained stable in TRP1-Fer but was downregulated in TRP4-Fer and EPG conditions. FoxO1 is critical in maintaining the balance between T cell activation and apoptosis.^45^

STING (Stimulator of Interferon Genes) is crucial for detecting cytosolic DNA and initiating the type I interferon response.^46^ The expression of STING was slightly downregulated in TRP1-Fer, but showed a moderate increase in EPG-expressing cells.

Furthermore, the expression of several critical genes involved in various immune and metabolic pathways was examined to further investigate the tools’ impact on the cells (Figure 4A).

Taken together, these results highlight the ability of magnetogenetic tools to fine-tune T cell responses, offering insights into their potential application in immunotherapy and immune modulation. The observed variations among different magnetogenetic tools may stem from the fact that each tool likely has its own optimal conditions for induction. Factors such as the strength and frequency of the magnetic field, the duration of exposure, and the specific molecular mechanisms targeted by each tool can all influence their effectiveness. Additionally, the unique properties of each magnetogenetic construct, such as its sensitivity to magnetic fields or its interaction with intracellular signaling pathways, might require different induction protocols to achieve maximal or desired effects.

The magnetogenetic tools indicated that the induction system alone could lead to a deactivation or suppression of T cell function, as shown by the downregulation of key activation markers such as CD69, IL2, and IL6 (Figure 3C). This deactivation may be due to a protective feedback mechanism, where the cells reduce their activity to prevent over-stimulation from the initial calcium influx induced by the system. However, when antigenic stimulation (via anti-CD3 and anti-CD28 antibodies) was introduced alongside the magnetogenetic induction, T cell activation was amplified, as evidenced by the increased expression of activation markers and cytokines (Figure 4A). This suggests that the magnetogenetic tools can potentiate T cell activation when combined with external stimulatory signals, overcoming the deactivation phase and driving a stronger immune response. This dual-phase effect could be leveraged to control T cell activation precisely, allowing for modulation between resting states and active immune responses based on the presence of antigens (Figure 4B). This highlights the potential of magnetogenetic tools to not only regulate T cell deactivation but also to enhance T cell activation when coupled with the right external stimuli.

### Proteomics Analysis of the Tools’ Effect

To further investigate the changes in Jurkat cells induced by magnetic field exposure without antigen stimulation, we analyzed their protein profile using proteomics. We examined cells at 1, 2, and 6-hour induction time points. The 6-hour induction results showed the most significant changes across a majority of proteins, prompting us to select this time point for more detailed analysis. Our findings confirmed that both the EPG and engineered TRPV ion channels were capable of altering cell function. To further clarify these changes, we categorized key proteins based on their functions.

### T Cells Specific Proteins

The acquired data showed a significant downregulation of critical T cell proteins such as CD1 and CD4, which are vital for T cell activation, antigen recognition, and differentiation in non-stimulated cells (Figure 5 and see also the supplementary information). This suggests that non-stimulated magnetogenetic induction may impair basic T cell functions. Notably, CD69 and IL2, key activation markers,^19^ were upregulated in stimulated cells, suggesting that antigenic stimulation alongside magnetic induction enhances cell activation despite downregulation seen in non-stimulated conditions.

**Figure 5.**
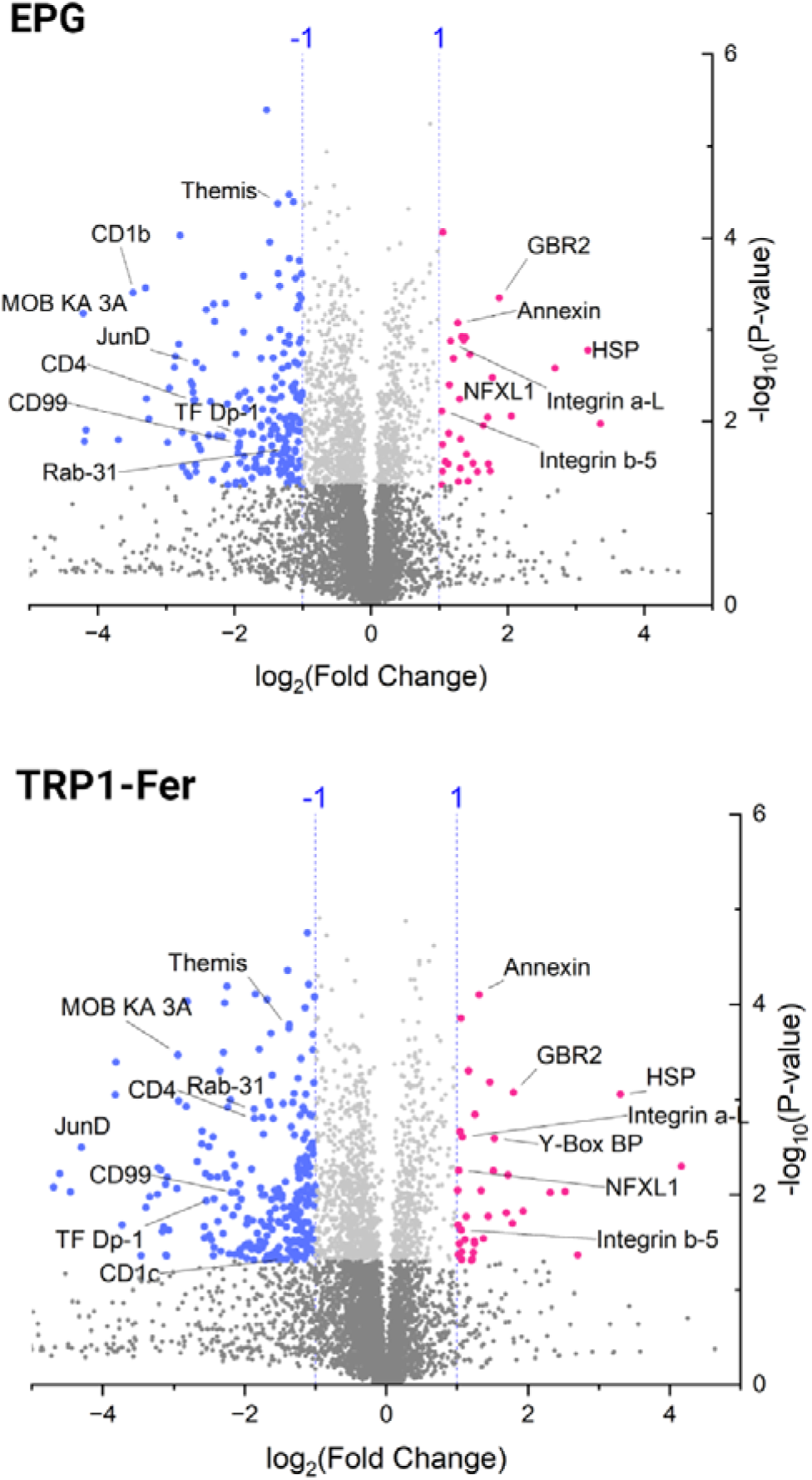
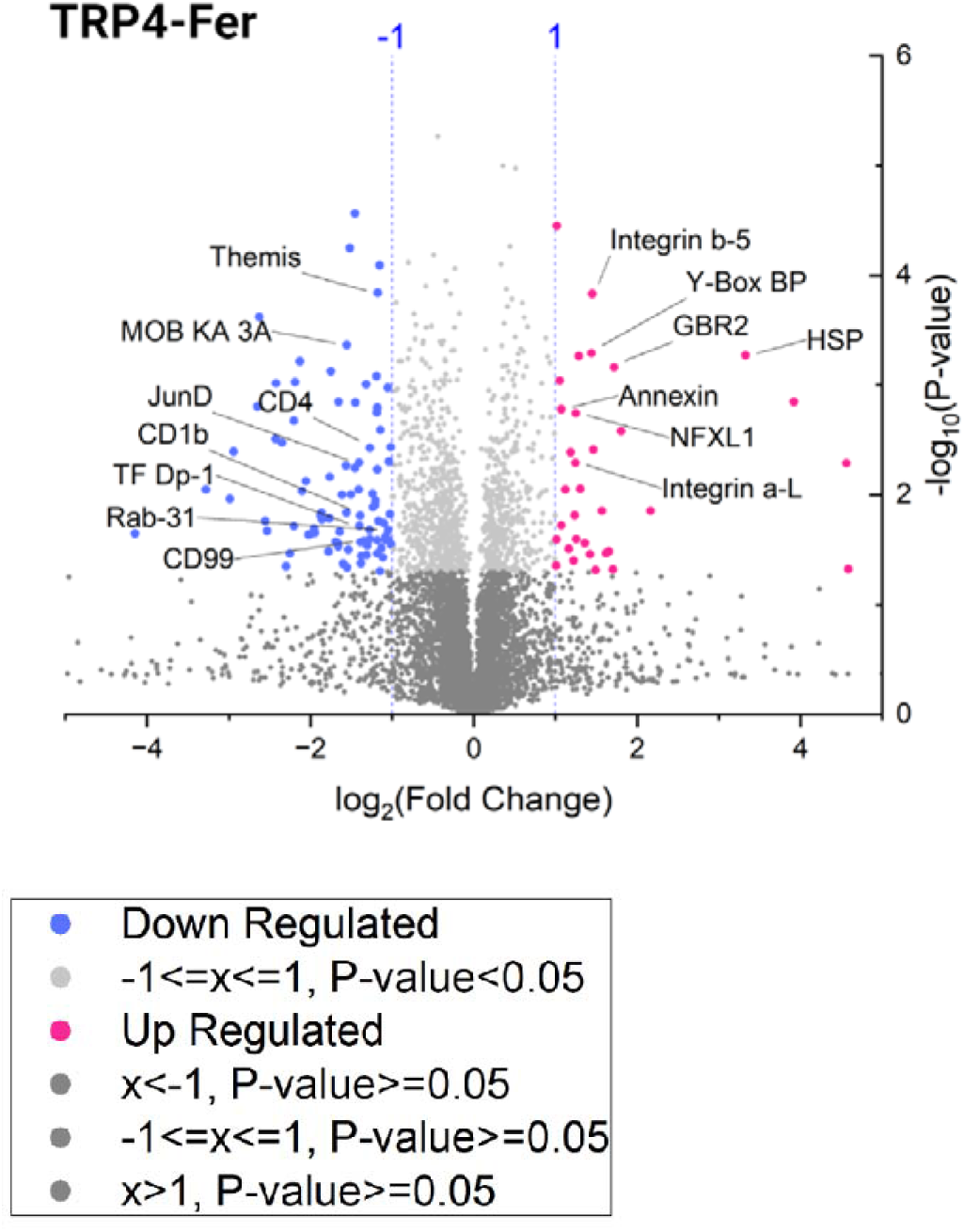
Proteomics analysis. Volcano plots of proteomics data showing the increase and decrease of T cells specific proteins after 6 hours of magnetic induction in Jurkat cells without antigen stimulation.

The proteomics data also showed decreased expression of CD99 and Rab-31, proteins involved in cell adhesion, migration, and vesicular trafficking.^47^ This downregulation implies that T cells might have reduced mobility and communication abilities in non-stimulated conditions, potentially hindering their ability to engage in immune surveillance and response. This aligns with the downregulation of CXCR6 in TRP4-Fer-expressing cells from the qPCR data, indicating impaired T cell migration in certain magnetogenetic conditions. However, CXCR3 and CXCR6 expression was stable or upregulated in EPG and TRP1-Fer expressing cells, suggesting that under stimulation, these cells may retain some ability to migrate effectively towards sites of inflammation.

The downregulation of other transcription factors like THEMIS and JunD, which are involved in T cell receptor (TCR) signaling and transcription regulation,^48,49^ further suggests that T cell activation and differentiation could be compromised.

Additionally, the observed downregulation of MOB kinase activator 3A, Transcription factor Dp-1, and F-box/LRR-repeat protein 6 reveals key insights into the changes within T cells induced by magnetogenetic tools. MOB kinase activator 3A, a regulator in the Hippo signaling pathway, suggests decreased control over cell proliferation and survival,^50^ potentially compromising T cell expansion and effectiveness. The downregulation of Transcription factor Dp-1 indicates a reduced readiness for cell division, which is essential for robust immune responses through rapid T cell proliferation.^51^ Meanwhile, the decrease in F-box/LRR-repeat protein 6 highlights impaired regulation of protein degradation, which is crucial for maintaining cellular balance by managing signaling. Notably, the expression of MOB kinase activator 3A was stable in the stimulated cells and the upregulation of IL2, a major driver of T cell proliferation, implied that antigenic stimulation could overcome this proliferation deficit in certain contexts. This suggests that while basal T cell proliferation might be compromised by magnetogenetic tools alone, external stimulation could restore and enhance this proliferative capacity.

Furthermore, the upregulation of several proteins in deactivated cells likely reflects compensatory or stress-related responses rather than active cell signaling. GRB2-related adapter protein 2 was elevated, suggesting an attempt to maintain signaling capacity,^52^ although this may not suffice for full cell activation. Similarly, the increase in Annexin A1, a protein involved in anti-inflammatory processes, points to a shift toward a regulatory or deactivated state.^53^ The rise in Heat shock protein 105 kDa indicates a cellular stress response,^54^ as this protein plays a key role in protecting cells from stress-induced damage by stabilizing proteins. Finally, the upregulation of NF-X1-type zinc finger protein NFXL1 could reflect enhanced transcriptional activity focused on managing cellular stress, repair, and survival mechanisms, rather than activation. Together, these changes suggest that while the cells are deactivated, they are engaging in protective mechanisms to mitigate the effects of stress induced by magnetogenetic tools.

### Channels and Ions-Related Proteins

Due to the effects of magnetogenetic tools on calcium concentrations and the associated impacts on ion-related proteins, we analyzed the involved channels and proteins (Figure 6A and see the supplementary information). The induction of magnetogenetic tools in cells appears to have led to a significant downregulation of calcium-related signaling pathways in non-stimulated cells by antigens, particularly through the suppression of ion channels and associated proteins.

**Figure 6.**
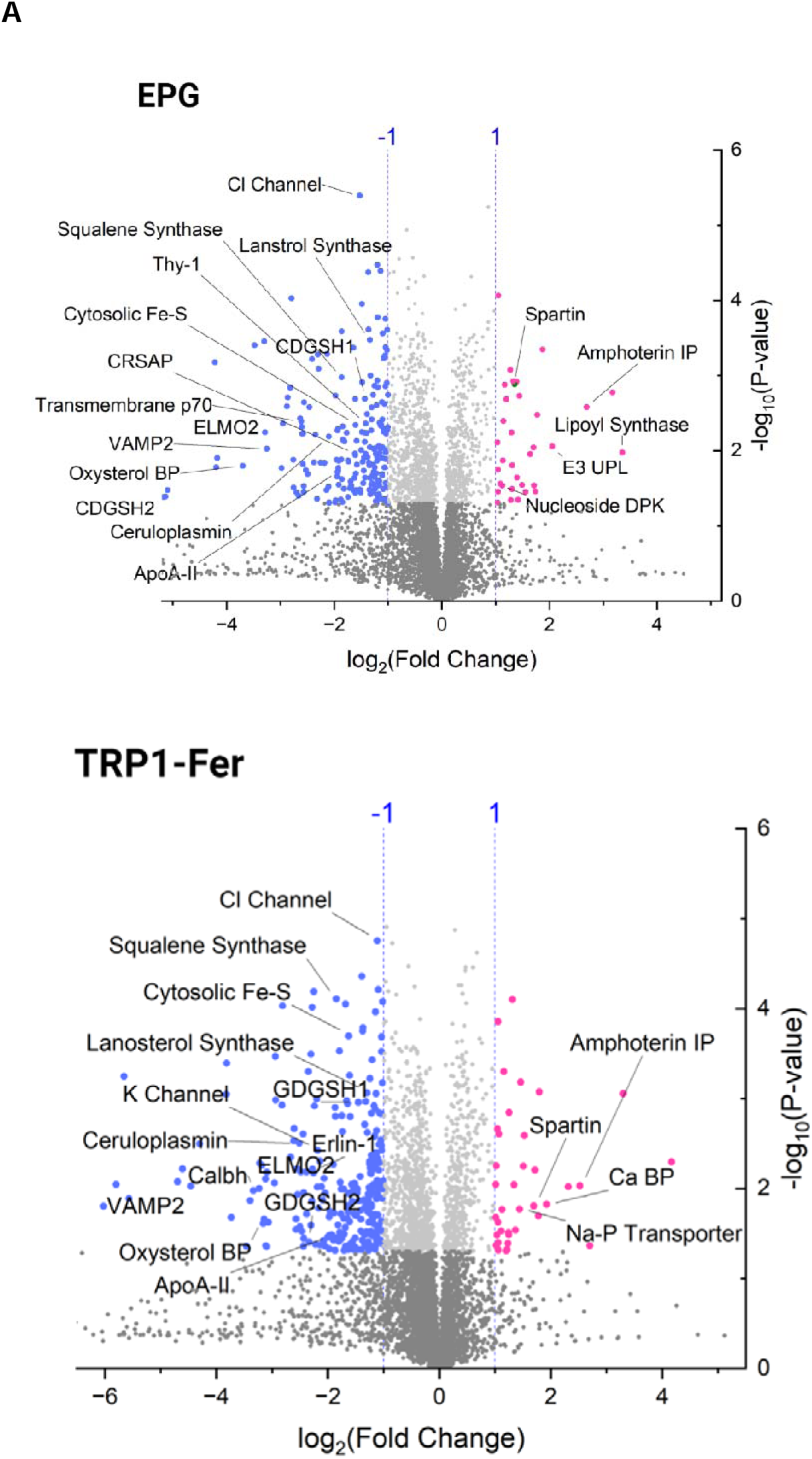
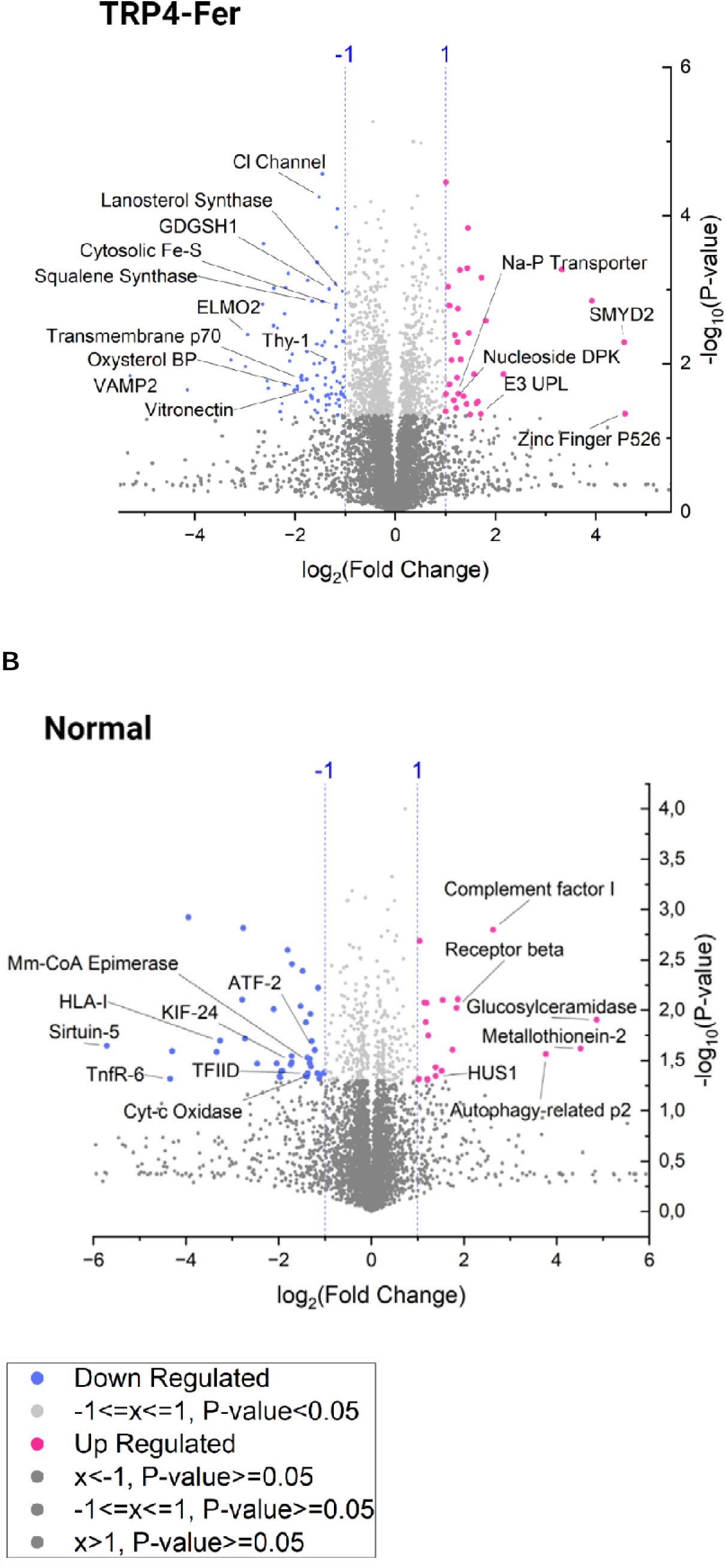
Proteomics analysis. (A) Volcano plots of proteomics data showing the increase and decrease of various proteins after 6 hours of magnetic induction in Jurkat cells expressing different tools without antigen stimulation. (B) Proteome profile of normal Jurkat cells without expressing tools and antigens after 6 hours magnetic induction.

The downregulation of the Calmodulin-regulated spectrin-associated protein underscores a reduction in calcium-dependent processes. This protein is essential for cytoskeletal stability and cellular motility, and its decrease suggests that the induced T cells may experience reduced structural changes, potentially limiting their ability to migrate and interact with other cells.^55,56^ However, the qPCR data of stimulated and magnetically induced cells showed a significant upregulation of **Calcineurin B Homologous Protein 3 (Calbh)** in both TRP1-Fer and EPG-expressing cells and the proteomics data of antigen-stimulated cells, indicating heightened calcium-mediated signaling under these conditions.^27^

The reduction in channels such as the Mitochondrial Potassium Channel and Chloride Intracellular Channel Protein 1 further indicates that these cells might be less equipped to adjust their ionic balances to accommodate the reduced calcium signaling.

Conversely, certain proteins, like the Sodium-Dependent Phosphate Transporter 1, showed an increase in expression, which could suggest a shift in cellular priorities towards different metabolic processes. The observed regulation of various calcium-handling proteins and channels, suggests a complex and somewhat maladaptive response by the cells to the challenges posed by magnetogenetic tool induction. This response likely reflects an effort to stabilize ion homeostasis and maintain cellular integrity under stress but may also indicate an inability to fully compensate for the reduced calcium signaling and metabolic activity.

### Cell Membrane-Related Proteins

The induction of magnetogenetic tools in T cells, while providing advanced capabilities for manipulating cellular functions, also presents potential risks to the integrity of the cell membrane (Figure 6A and see also the supplementary information). The cell membrane is crucial for maintaining cellular homeostasis, mediating cell signaling, and ensuring proper interaction with the extracellular environment. However, the generation of reactive oxygen species (ROS) and heat during the application of these tools can lead to membrane damage, compromising these vital functions.^57^ The differential expression of various membrane-related proteins in this context offers insights into both protective adaptations and potential vulnerabilities of Jurkat cells.

In non-stimulated cells, downregulation of proteins such as Oxysterol-binding protein-related protein 5, Lanosterol synthase, Vesicle-associated membrane protein 2, Transmembrane protein 135, Transmembrane protein 70 (mitochondrial), Thy-1 membrane glycoprotein, FAS-associated death domain protein, Squalene synthase, and Apolipoprotein A-II was observed (Figure 6A and the supplementary information). These proteins are involved in lipid transport, membrane repair, and cell adhesion, and their decreased expression suggests that T cells may struggle to maintain membrane integrity and fluidity. For example, the reduction in Oxysterol-binding protein-related protein 5 and Lanosterol synthase indicates potential challenges in maintaining lipid distribution and cholesterol biosynthesis, which are critical for membrane stability.^58,59^

The downregulation of Vesicle-associated membrane protein 2 (VAMP2) and Transmembrane proteins 135 and 70 suggests a reduced capacity for vesicular trafficking and membrane repair.^59,60^ This could make T cells more susceptible to damage, as the efficiency of membrane repair mechanisms is compromised. Similarly, the decreased expression of Thy-1 membrane glycoprotein may impair cell adhesion and immune synapse formation,^61^ which are crucial for effective immune responses.

### Mitochondria-Related Proteins

The proteomics analysis revealed significant downregulation of key mitochondrial proteins in Jurkat cells subjected to magnetogenetic induction, suggesting compromised energy production and metabolic flexibility. Proteins such as Succinate dehydrogenase cytochrome b560 subunit and Succinate dehydrogenase assembly factor 2, both crucial components of the electron transport chain and citric acid cycle,^62^ were significantly downregulated. This reduction points to impaired mitochondrial respiration and ATP synthesis, which are essential for the energy demands of activated T cells. Also, the downregulation of Mitochondrial carnitine/acylcarnitine carrier protein, involved in transporting fatty acids into the mitochondria for beta-oxidation, suggests a reduced capacity for utilizing fatty acids as an alternative energy source.^63^ This could limit the metabolic flexibility of T cells, particularly in conditions where glucose is scarce or during prolonged periods of activation (see the supplementary information).

Moreover, the proteomics data demonstrated a decrease in iron-sulfur cluster assembly proteins like CDGSH1, CDGSH2, and cytosolic iron-sulfur components, which are essential for mitochondrial function.^64,65^ The downregulation of these proteins suggests compromised mitochondrial activity, which is crucial for supporting the energy-intensive processes of activated T cells (Figure 6A). However, the stimulated cells data did not show any downregulation of these genes.

Furthermore, Ceruloplasmin, another protein related to iron (Fe) homeostasis, in non-stimulated cells also showed a decrease in expression. This copper-carrying protein plays a crucial role in iron metabolism by oxidizing Fe²⁺ (ferrous iron) to Fe³⁺ (ferric iron), facilitating its incorporation into transferrin for transport. The reduced levels of Ceruloplasmin could disrupt iron balance within the cells, potentially leading to the accumulation of free iron, which can catalyze the production of harmful ROS (Figure 6A).^66^

### Other Proteins

The induction of magnetogenetic tools in Jurkat cells led to significant downregulation of proteins related to lipid metabolism, cellular motility, and extracellular matrix interactions. Proteins like Erlin-1, Apolipoprotein A-II (ApoA-II), Engulfment and Cell Motility Protein 2 (ELMO2), and Vitronectin were notably reduced, indicating several challenges in cell functionality. Erlin-1 is involved in the ER-associated degradation (ERAD) pathway, which manages protein quality control.^67^ Its downregulation suggests that magnetogenetic tools might impair the ability of T cells to cope with stress, potentially leading to ER stress and dysfunction.

Apolipoprotein A-II (ApoA-II) plays a key role in lipid metabolism and stabilizing high-density lipoproteins (HDL). The observed decrease in ApoA-II suggests a potential reduction in lipid metabolism, which could hinder membrane synthesis and energy production, thereby limiting the ability of T cells to maintain membrane fluidity and meet the energetic demands of activation and proliferation.^68^

ELMO2 was also downregulated, indicating potential impairments in immune surveillance. Similarly, Vitronectin, a glycoprotein crucial for cell adhesion and complement system regulation, was reduced. This suggests that T cells may struggle to adhere to and migrate through the extracellular matrix (ECM), possibly weakening immune cell trafficking and wound healing responses.^69^

### Magnetic field effects on normal Jurkat cells

In this study, the proteomic analysis of normal Jurkat cells (without magnetogenetic tools) exposed to a magnetic field demonstrated changes in the expression of various proteins. These changes suggest that Jurkat cells are dynamically responding to magnetic field exposure, which affects their structural integrity, transcriptional regulation, and overall metabolic capacity (Figure 6B and see the supplementary information).

In mitochondrial and metabolic proteins, Key metabolic proteins, such as Cytochrome c oxidase subunit 7A-related protein, Methylmalonyl-CoA epimerase, and NAD-dependent protein deacylase sirtuin-5, showed significant downregulation. These proteins are vital for maintaining mitochondrial function, energy production, and metabolic homeostasis. Their decreased expression suggests that cells may face challenges in maintaining energy production and structural integrity under stress, potentially leading to a compromised immune response.^70,71^

Proteins involved in maintaining cytoskeletal structure, such as MAP/microtubule affinity-regulating kinase 3 and Kinesin-like protein KIF24, also showed reduced expression. The downregulation of these proteins could impair intracellular transport and structural stability, critical for cell motility and function.^72,73^

Transcriptional regulators like Cyclic AMP-dependent transcription factor ATF-2 and Transcription initiation factor TFIID subunit 3 were also downregulated, indicating potential disruptions in the transcriptional regulation necessary for T cell survival and stress response.^74,75^ These reductions may compromise the cells’ ability to regulate gene expression effectively, hindering their adaptability to environmental stress.

Among DNA damage response and repair, despite these challenges, certain proteins showed increased expression, possibly as a compensatory mechanism. Checkpoint protein HUS1, involved in the DNA damage response, and Histone H2A.Z, associated with chromatin stability, were upregulated. This suggests that cells are enhancing their ability to repair DNA and maintain genomic stability under magnetic field exposure.^76,77^

Proteins involved in managing oxidative stress, such as Metallothionein-2 and Lysosomal acid glucosylceramidase, were significantly upregulated, indicating enhanced protection against oxidative damage. The upregulation of these proteins suggests that cells may be attempting to bolster their antioxidant defenses and autophagic capacity, essential for cellular homeostasis.^78,79^

Additionally, proteins critical for immune signaling and activation, such as cell receptor beta constant 1 and Complement factor I, showed increased expression. This suggests that cells might be enhancing their immune responsiveness and capacity to regulate the complement system, potentially improving their ability to respond to pathogens.^80,81^

Exposure to magnetic fields induces a complex response in Jurkat cells, characterized by the downregulation of key mitochondrial, structural, and transcriptional proteins, which may compromise their function under stress. However, the upregulation of proteins involved in DNA repair, oxidative stress response, and immune activation indicates that cells are simultaneously activating compensatory mechanisms to mitigate damage and maintain functionality. These findings highlight the importance of further research into the effects of magnetic fields on T cell biology, particularly in therapeutic contexts where T cell efficacy is critical.

### Confirmation of the System’s Effect on T Cells

To explore whether the effects of magnetogenetic tools in primary T cells mirror those observed in Jurkat cells, we isolated T cells from human peripheral blood mononuclear cells (PBMC) samples and transduced them with our custom-made lentiviruses. The cells were then exposed to a magnetic field, both with and without antigenic stimulation, and the expression of activation markers CD69 and IL2 was measured to assess T cell activation. Our results demonstrated that the T cells responded similarly to Jurkat cells, showing comparable patterns of activation and deactivation (Figure 7). This experiment confirms that the underlying mechanisms driving activation in Jurkat cells are consistent with those in T cells isolated from PBMCs, validating the relevance of Jurkat cell models for studying the effects of magnetogenetic tools.

**Figure 7.**
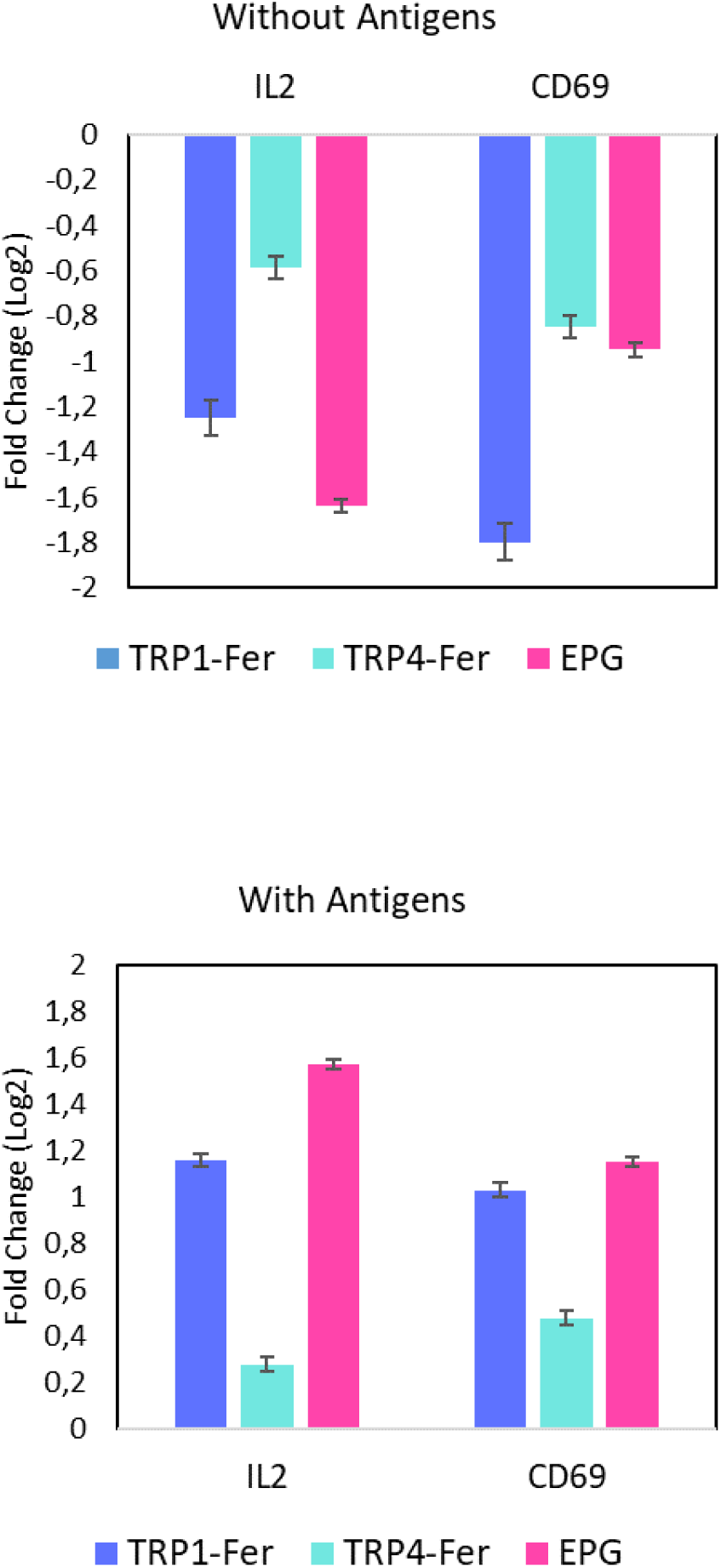
Expression of CD69 and IL2 genes in T cells after 6 hours of magnetic induction with and without adding anti-CD3 and anti-CD28 antibodies.

## Conclusion

This study demonstrated the potential of magnetogenetic tools, specifically TRP1-Fer, TRP4-Fer, and EPG, to modulate Jurkat and T cell behavior through magnetic field induction, influencing calcium signaling and gene expression. While magnetic induction alone led to cell deactivation, the combination of magnetic induction with antigenic stimulation significantly amplified cell activation compared to controls. Notably, the different magnetogenetic tools exhibited varying strengths and functions in our magnetic induction setup, with each tool showing distinct (but similar) effects on cell activation and deactivation.

The proteomics and gene expression analysis revealed adaptive cellular responses, including stress-related changes and modulation of signaling pathways, aimed at maintaining cellular homeostasis despite deactivation. These findings highlight the delicate balance between the deactivating effects of magnetic induction and the potential for antigenic stimulation to enhance cell responsiveness, with the choice of magnetogenetic tool playing a crucial role.

Overall, this research emphasizes the value of magnetogenetic tools in non-invasively controlling immune cell function. With further optimization, these tools hold great promise for applications in immunotherapy and synthetic biology, where precise control over immune responses is key to therapeutic success.

## Material and Methods

### Plasmids construct and the cell lines

In this study, 3 different tools were investigated. TRPV1 and TRPV4 channels as well as EPG protein domain. To make the lentiviruses, these 3 genes were synthesized and were cloned into S/MAR-Minicircle plasmids to generate McTFer1, McTFer4, and McEPG. To track the transfected cells, expression confirmation, and sorting the cells by Flowcytometry, GFP marker for McTFer1, and mCehrry for McTFer4 and McEPG plasmids were used. Then,

### Strains and plasmid preparation

For cloning and plasmid amplification, *E. coli* DH5α was utilized. Prior to transformation, the cells were incubated overnight at 37°C and 250 rpm and then rendered chemically competent with CaCl_2_. Single colonies were selected and cultured in LB medium supplemented with either 50 µg/mL kanamycin or 100 µg/mL ampicillin at 37°C and 250 rpm.

To assess the system in Hek-293T cells, genes were inserted into S/MAR minicircle plasmids. To generate the MC TRP1-Fer, MC TRP4-Fer, and MC EPG plasmids, distinct genes were cloned and inserted into S/MAR minicircle plasmids. Modifications were achieved by amplifying and linearizing the plasmid using primers with corresponding sequences and overhangs, employing the GeneArt™ Gibson Assembly HiFi Master Mix (Thermo Fischer Scientific). Phusion Hot Start II High-Fidelity PCR Master Mix (Thermo Fischer Scientific) was used for PCR reactions.

For plasmid extraction, the NucleoSpin Plasmid Mini kit (MACHEREY-NAGEL) was used for minipreparations, while the QIAGEN Plasmid Plus Maxi kit was used for maxipreparations. The DNA concentration was determined using a NanoDrop™ 2000/2000c spectrophotometer (Thermo Scientific™). To confirm the DNA sequence of all plasmid constructs, samples were prepared accordingly and sent to Eurofins Scientific for sequencing utilizing the Mix2Seq Kit NightXpress.

### S/MAR Minicircle plasmid preparation

Minicircles were generated from the full-length parental plasmid using PhiC31 integrase, which enabled recombination between the PhiC31 attB and attP sites present on the parental plasmid. This process produced two distinct products: the minicircle, which lacked any bacterial backbone DNA, and the remaining portion of the parental plasmid. To remove the bacterial backbone-containing plasmid in *E. coli*, the I-SceI endonuclease was used to recognize and cleave the I-SceI sites, leading to its degradation. For effective minicircle production, the *E. coli* strain ZYCY10P3S2T was employed, which has an arabinose-inducible system for simultaneous expression of both PhiC31 integrase and I-SceI endonuclease. Additionally, this strain harbors a modified arabinose transporter, the LacY A177C gene. By adding arabinose to the media, the expression of the integrase and endonuclease was induced, enabling the separation of the parental plasmid into minicircles and the plasmid containing the bacterial backbone.

For the generation of minicircle plasmids, we used the MC-Easy™ Minicircle DNA Production Kit (System Biosciences, SBI). The procedure began by inoculating 2 mL of LB-Kanamycin medium with the ZYCY10P3S2T strain transformed with the minicircle plasmid, followed by incubation for 1 hour at 30°C with shaking at 250 rpm. This was then transferred to 200 mL of LB-Kanamycin medium and incubated for 16 hours at 30°C with shaking. Afterward, 200 mL of induction medium containing arabinose was added, and the culture was incubated for another 3 hours at 30°C. The temperature was then raised to 37°C for 1 additional hour before the cells were harvested for mini/maxiprep plasmid extraction.

To confirm the successful removal of the bacterial backbone, the size and structure of the purified plasmids were verified via restriction enzyme digestion, followed by analysis using agarose gel electrophoresis.

### Cell culture and gene delivery to mammalian cells

Normal HEK293T (Hek) cells, obtained from the American Type Culture Collection (ATCC), were cultured at 37°C with 5% CO2 in DMEM supplemented with 10% fetal bovine serum (FBS) and 1% PenStrep (100 U/mL penicillin and 100 μg/mL streptomycin). The Jurkat cell line, obtained from ATCC (Clone E6-1), was cultured under the same conditions but in RPMI medium. Prior to the experiments, all cell cultures were tested for mycoplasma contamination.

For delivering plasmids to Hek cells, both lipofection technique was employed. Jurkat cells were exclusively transfected via electroporation and lentiviral transduction.

All DNA plasmids were transfected into Hek cells using Lipofectamine™ LTX Reagent (Thermo Fisher Scientific). Cells were seeded 24 hours prior to transfection to ensure high viability and suitable confluency. Subsequently, the DNA-lipid complex was added to the cells and incubated at 37°C with 5% CO_2_ for 3-6 days.

For electroporation of Jurkat cells, the Lonza™ P3 Primary Cell 96-well Nucleofector™ Kit (Lonza) was utilized. Briefly, after the cells were washed with PBS buffer, they were suspended in P3 Primary Cell Nucleofector™ Solution (Lonza) and mixed with the desired amount of plasmid per reaction (ranging from 0.2 to 5 μg). This cell-plasmid mixture was then loaded into the electroporation wells. Electroporation was carried out using the Jurkat CL120 program on a 4D-Nucleofector® Core Unit (Lonza) machine. Following electroporation, additional media was added to the cells, which were subsequently incubated at 37°C with 5% CO_2_ for 3-5 days.

Notably, in this study, 150,000 cells were used for each reaction for both cell lines. Different plasmid ratios and concentrations (ranging from 0.5 µg to 2.5 µg) were tested.

Jurkat cell activation was induced using Human (0.5 µg/ml) T-Activator anti-CD3 and anti-CD28 antibodies (BioLegend) for cell expansion and activation.

### Lentivirus production and T cells transduction

Lentiviral particles were produced by transfection of a 3^rd^ generation lentivirus system (pMD2.G was a gift from Didier Trono (Addgene plasmid 12259 ; http://n2t.net/addgene:12259 ; RRID:Addgene_12259), [1] pMDLg/pRRE was a gift from Didier Trono (Addgene plasmid 12251 ; http://n2t.net/addgene:12251 ; RRID:Addgene_12251), and [2] pRSV-Rev was a gift from Didier Trono (Addgene plasmid 12253 ; http://n2t.net/addgene:12253 ; RRID:Addgene_12253)) altogether with transfer vector into Hek cells delivered by lipofectamine 3000 reagent. 24- and 48-hours post transfection, supernatants were harvested and mixed 1 to 3 with Lenti-X concentrator (Takara, 631232), cooled down to 4C for 30 minutes and spun down for 45 minutes at 4C.

Following centrifugation, supernatant was discarded, and the virus-containing pellet was resuspended in Opti-MEM (Gibco, 31985070) to obtain a highly concentrated virus solution, which was directly stored in 50ul aliquots at -80C until further use.

To transduce T cells, frozen PBMC samples were thawed, and a negative CD3 enrichment kit (StemCell, 19051) was used to isolate the T cell fraction. Following, cells were activated for 24 hours by using CD3/CD28 Dynabeads (ThermoFisher, 11132D) at a ratio of 3:1 bead/cell and 20 IU of IL-2 (Preprotech, 200-02) in X-Vivo media (Lonza, 02-053Q) supplemented with 5% human serum and Pen/Strep. The day after, T cells were transduced with variable amounts of the frozen virus and expanded for 9 more days, counting and subculturing every other day.

On day 10, cells were harvested, and beads were removed with a magnet. A small sample was used to assess transduction efficiency by flow cytometry and the rest were used fresh or cryopreserved for future use.

### Calcium concentration changes measurement

To measure calcium concentrations in T cells using the Fluo-4 Direct™ Calcium Assay Kit (Invitrogen), the following procedure was used. First, adherent T cells were cultured in 96-well microplates until near confluence. The Fluo-4 Direct™ calcium assay reagent was prepared by dissolving the components in the provided buffer according to the manufacturer’s instructions. For the assay, an equal volume of the 2X Fluo-4 Direct™ calcium reagent loading solution was added directly to the wells containing cells in culture medium, resulting in a 1X final concentration of the reagent. The plates were incubated at 37°C for 30 minutes followed by an additional 30 minutes at room temperature to ensure optimal dye loading. After the incubation, the fluorescence was measured using a microplate reader set to an excitation wavelength of 494 nm and an emission wavelength of 516 nm. The fluorescence intensity correlates with the intracellular calcium levels, providing a quantitative measure of calcium concentration changes in the cells.

### Experimental set-up for the magnetic system

To provide magnetic force for the induction, 5 permanent magnetic discs (Ф = 5 mm) were placed under the cell plate. The magnetic discs were spaced by 1 m to be at the center of the well (Figure S1). Eash magnetic disc provided a ∼ 50 μT of magnetic force measured at 1 mm distance from the surface of the disc. A mini-stepper motor (NEMA 11) with a linear stage is used to move the magnetic discs precisely under the wells. A pre-programmed code of the motor motion was uploaded into a microcontroller (Arduino Uno board), that was used to control the motor. A DC solenoid electromagnet (Molde no. MK 25, 6 Voltage and 0.5 Amper) was used for creating the alternating magnetic force (∼ 50 μT at the 1 mm discant from the surface of the electromagnet). The electromagnet was connected to the microcontroller to timely switch on/off the DC current, hence the magnet force. An NTA sensor (B57541G1103F005) was used to measure the temperature of medium in wells.

### Flow cytometry analysis

The cells were washed with PBS (Gibco) and resuspended in PBS. Cell viability staining was performed using Near-IR Live/Dead (NIR, Life Technologies, L10119) kit. Surface staining was carried out using a fluorochrome-labeled PE-conjugated anti-human CD11a (HI111) monoclonal antibody obtained from BioLegend.

The stained samples were analyzed on a Fortessa LSR flow cytometer (BD), and the data were analyzed using FlowJo V.10.9.0 software. Statistical significance was determined with a two-sided P value <0.05 considered statistically significant.

### Proteomics sample preparation

Briefly, cells were lysed using 50 uL of lysis buffer (consisting of 6M Guanidinium Hydrochloride, 10 mM TCEP, 40 mM CAA, 50 mM HEPES pH8.5). Samples were boiled at 95°C for 5 minutes, after which they were sonicated on high for 5x 60 seconds on and 30 seconds off in a Bioruptor Pico sonication water bath (Diagenode) at 4°C. 25 µL of sample were diluted 1:3 with digestion buffer (10% Acetonitrile, 50 mM HEPES pH 8.5) and 400 ng LysC (MS grade, Wako) was added in a 1:50 (enzyme to protein) ratio, and samples were incubated at 37°C for 4hrs. Samples were further diluted to a final 1:10 with digestion buffer and trypsin (MS grade, Sigma) was added in a 1:100 (enzyme to protein) ratio after which samples were incubated overnight at 37°C. Samples were acidified by adding 2% trifluoroacetic acid (TFA) to a final concentration of 1%. The peptides were desalted on a SOLAµ SPE plate (HRP, Thermo) following the same procedure as previously. Dried peptides were reconstituted in 12µL 2% ACN, 1%TFA.

### MS analysis

Peptides were loaded onto a 2cm C18 trap column (ThermoFisher 164946), connected in-line to a 15cm C18 reverse-phase analytical column (Thermo EasySpray ES904) using 100% Buffer A (0.1% Formic acid in water) at 750bar, using the Thermo EasyLC 1200 HPLC system, and the column oven operating at 30°C. Peptides were eluted over a 70 minute gradient ranging from 10% to 60% of Buffer B (80% acetonitrile, 0.1% formic acid) at 250 nl/min, and the Orbitrap Exploris instrument (Thermo Fisher Scientific) was run in DIA mode with FAIMS Pro^TM^ Interface (ThermoFisher Scientific) with CV of -45 V. Full MS spectra were collected at a resolution of 120,000, with an AGC target of 300% or maximum injection time set to ‘auto’ and a scan range of 400–1000 m/z. The MS2 spectra were obtained in DIA mode in the orbitrap operating at a resolution of 60.000, with an AGC target 1000% or maximum injection time set to ‘auto’, a normalised HCD collision energy of 32. The isolation window was set to 6 m/z with a 1 m/z overlap and window placement on. Each DIA experiment covered a range of 200 m/z resulting in three DIA experiments (400-600 m/z, 600-800 m/z and 800-1000 m/z). Between the DIA experiments a full MS scan is performed. MS performance was verified for consistency by running complex cell lysate quality control standards, and chromatography was monitored to check for reproducibility.

### Data analysis

The raw files were analyzed using Spectronaut^TM^ (version 18.6) spectra were matched against the human Uniprot database. The data is searched with the BGS workflow with Oxidation (M) and Acetyl on protein N-termini as dymanic modifications. Cysteine carbamidomethyl was set as a static modification. The proteotypicity filter wasturned off, MS quantity level to MS1 and local normalization was turned on.

### Gene expression analysis

RT-qPCR was used for gene expression. To this end, 200ng of total RNA were transcribed into cDNA using PrimeScript™ RT Master Mix (Takara). Each cDNA sample was quantified and diluted to 20ng/µl. For each cDNA sample, a qPCR master mix was prepared as follows: 10µl RealQ Plus 2x Master Mix Green Without ROX™ (Ampliqon), 1µl cDNA (20ng), and 8µl nuclease-free water; volumes were adjusted correspondingly to 50 qPCR reactions. For each sample, 19µl of the corresponding qPCR master mix were distributed in 48 wells within a 96-well PCR plate. 47 assay mixes were independently prepared (100µl forward primer 10µM + 100µl reverse primer 10µM, see Table S2), and 1µl of each assay mix was added to each one of the 48 reaction wells (final concentration 0.25µM). The assay mix for the S18 housekeeping gene was added in duplicate. The qPCR was performed as follows: 95°C for 15min; 40 cycles of 95°C for 20sec, 55°C for 30sec, and 72°C for 30sec, with fluorescent signal acquisition at the end of each cycle.

## Supplementary Information

**Figure S1.**
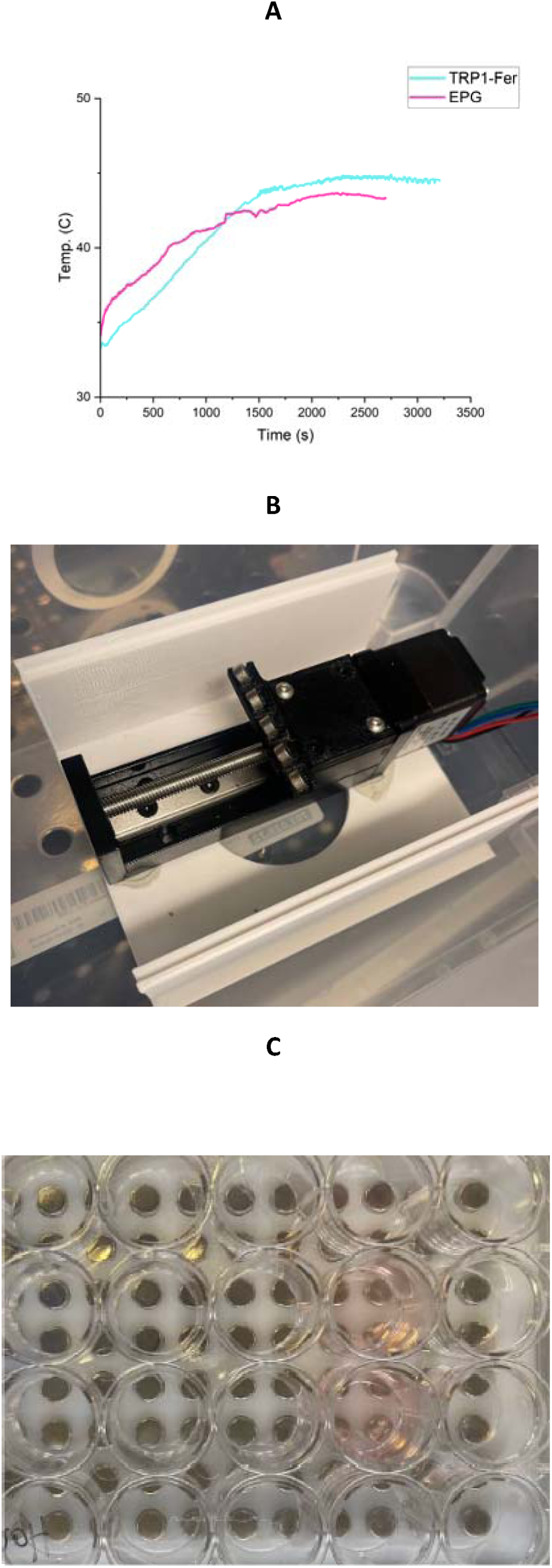
Advancement of magnetic induction system. (A) Temperature of the medium during alternating magnetic field (AMF) induction of TRP1-Fer and EPG-expressing Jurkat cells. (B) Design of the moving permanent magnets (MPM) setup. The cell plate is fixed, while a row of magnets moves beneath it, and (C) cells are shaken on a stationary magnet plate.

**Table S1.**
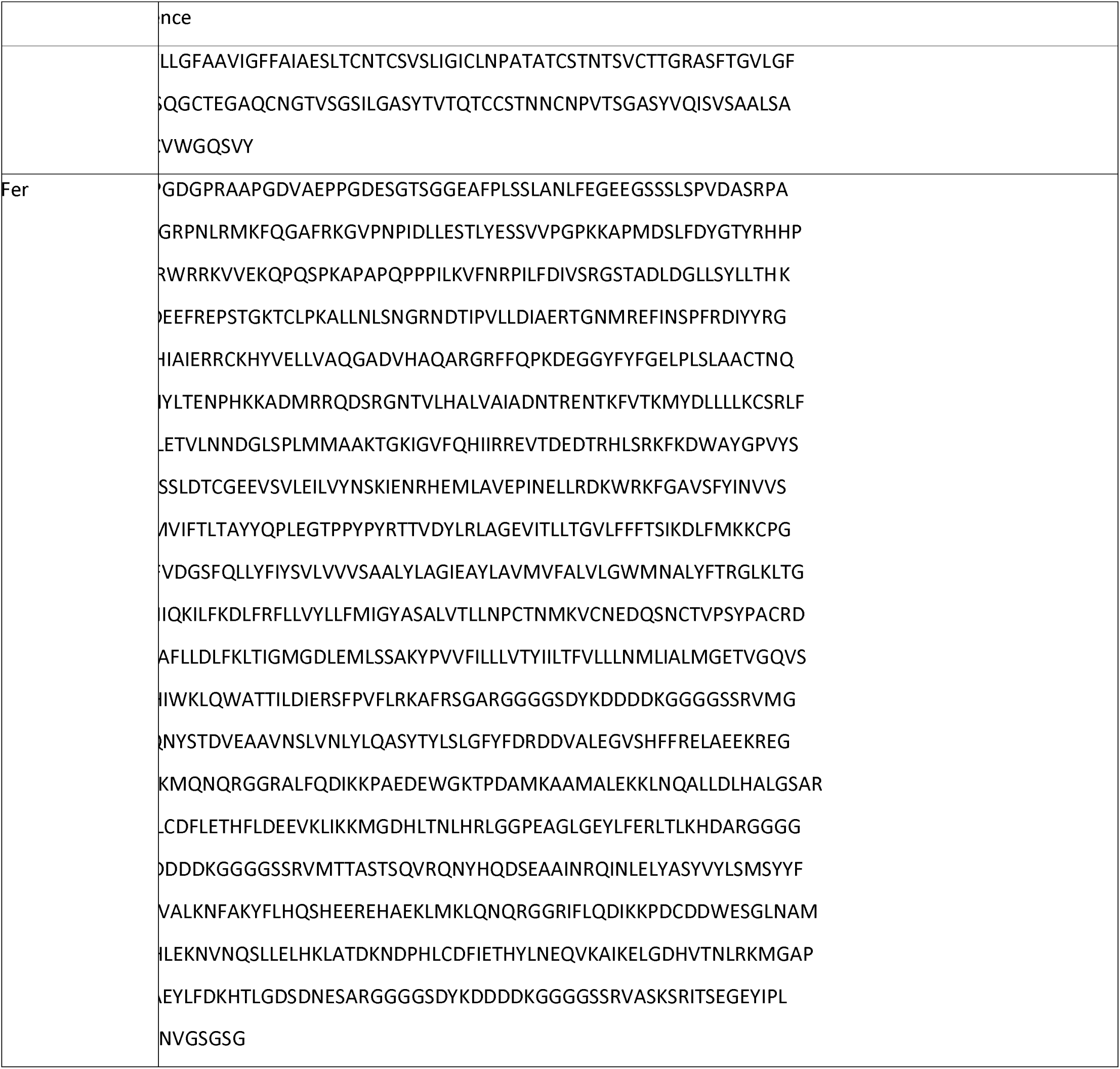
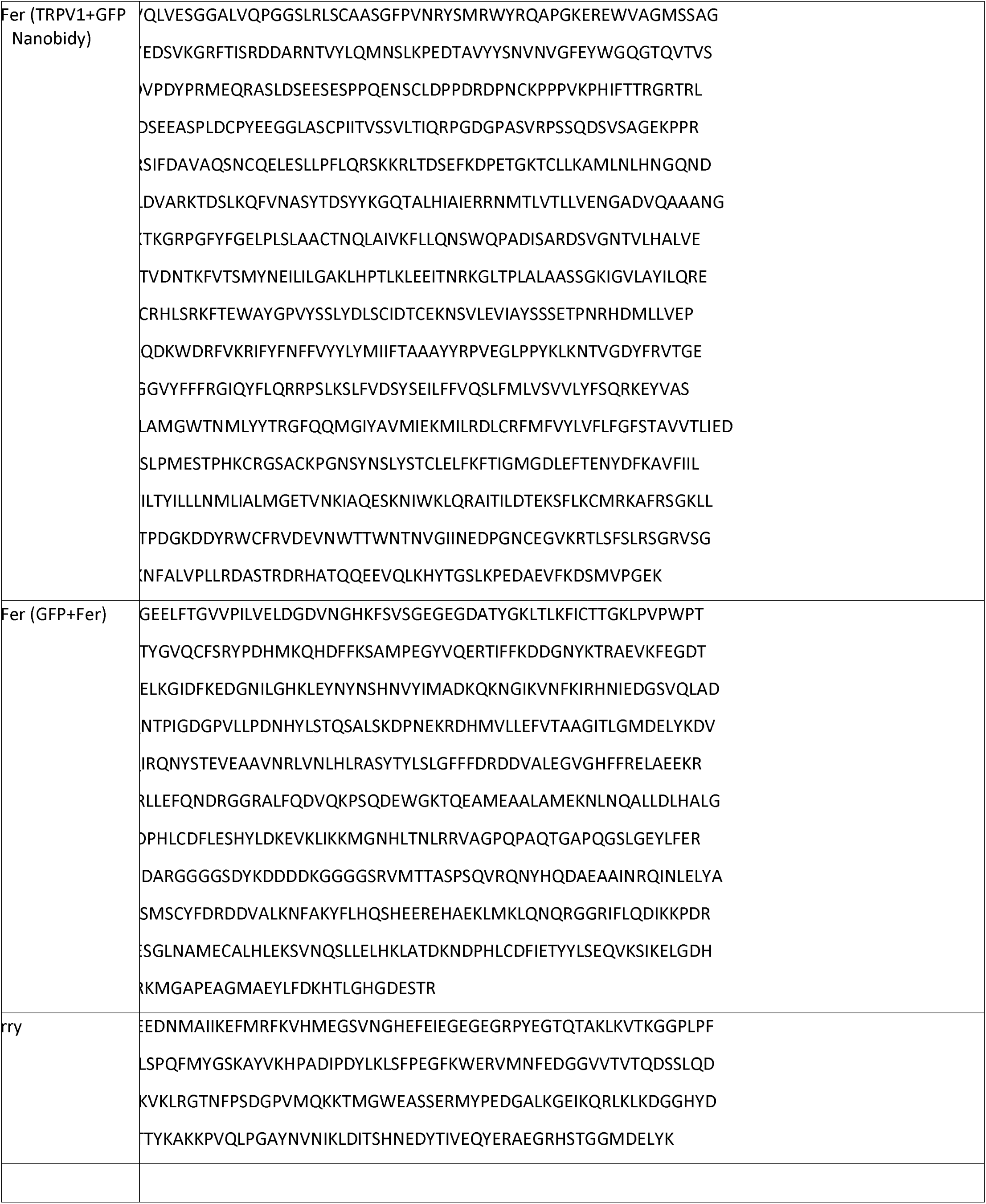
Protein sequences.

**Table S2.**
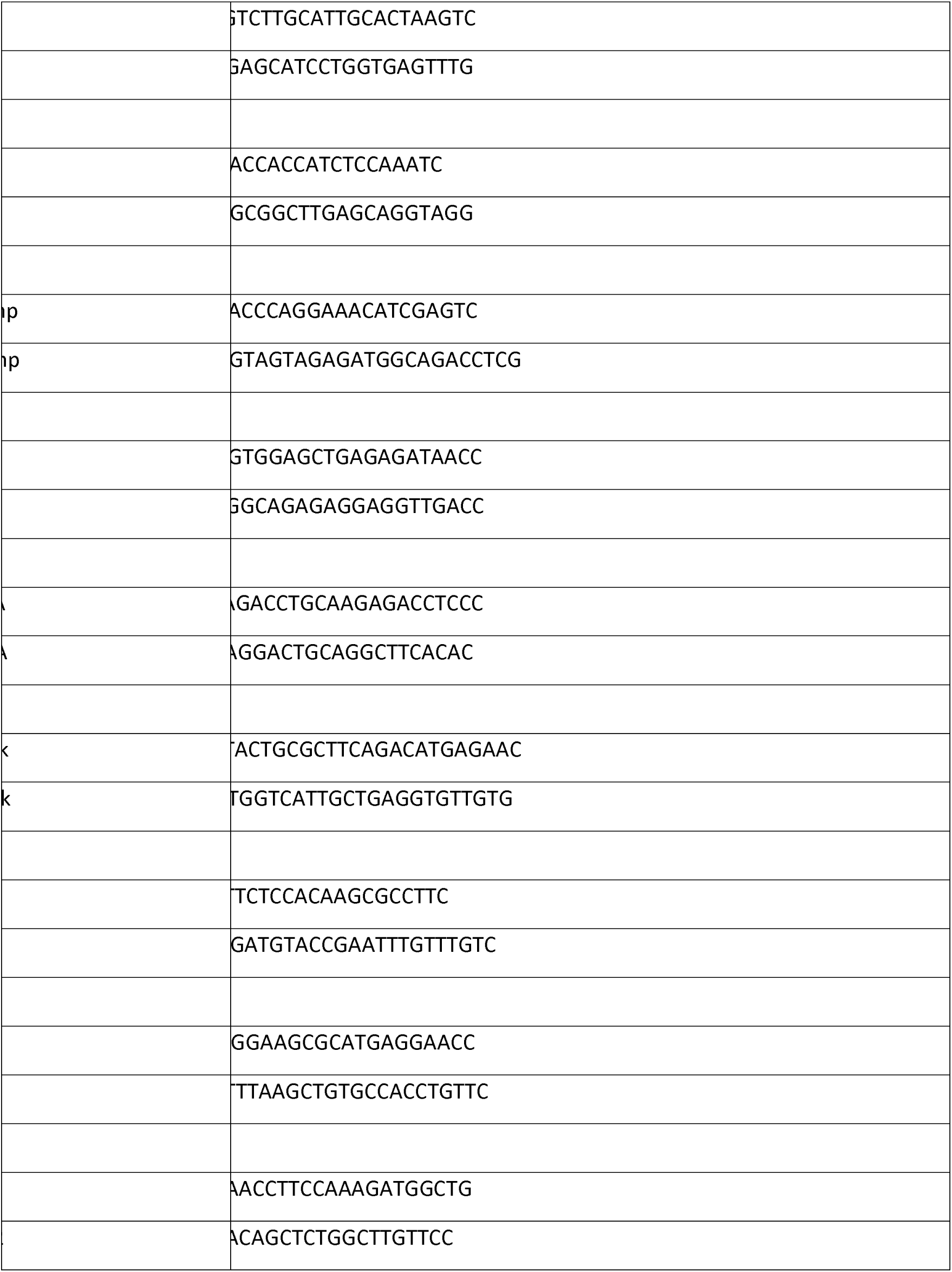
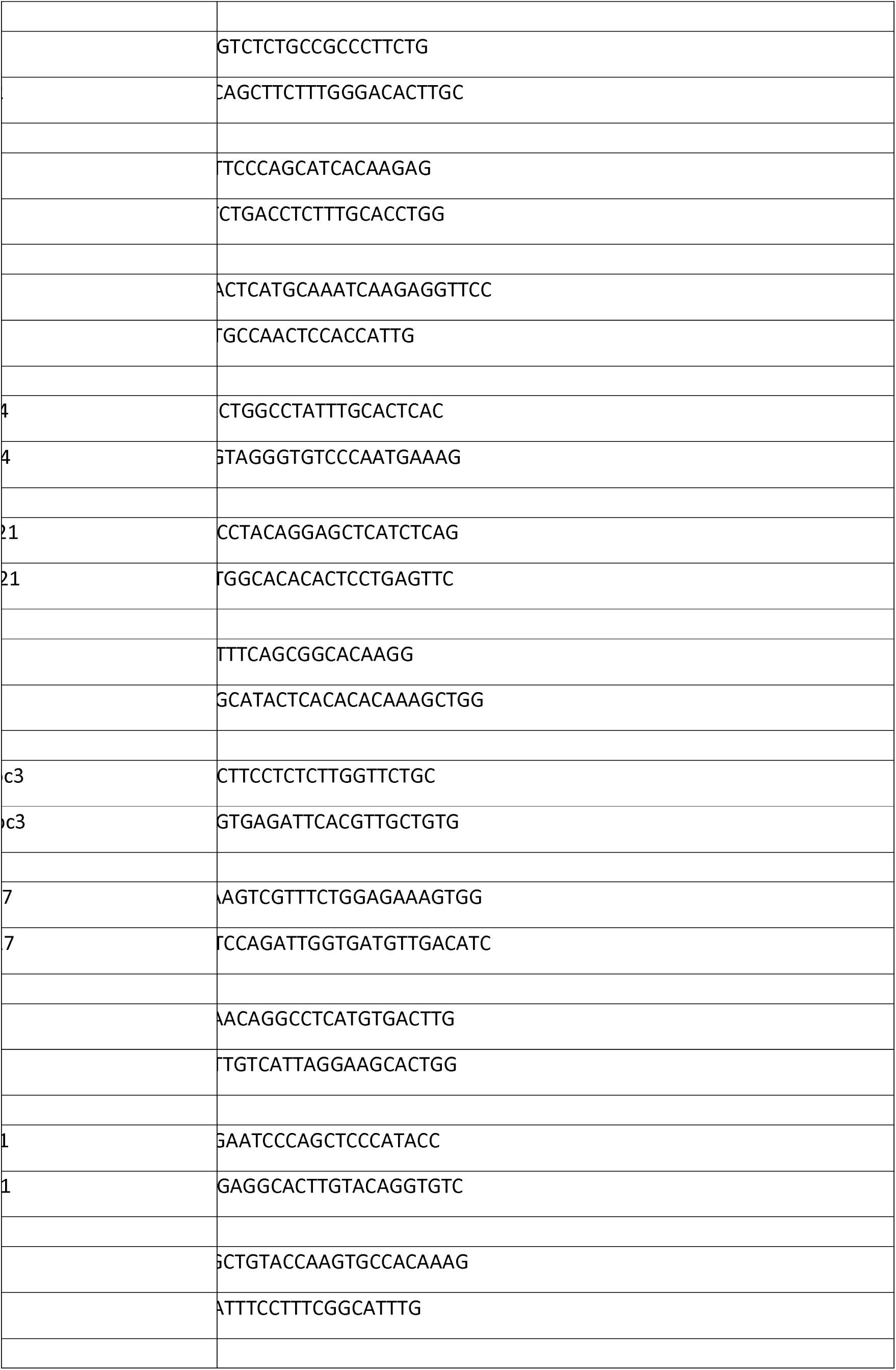
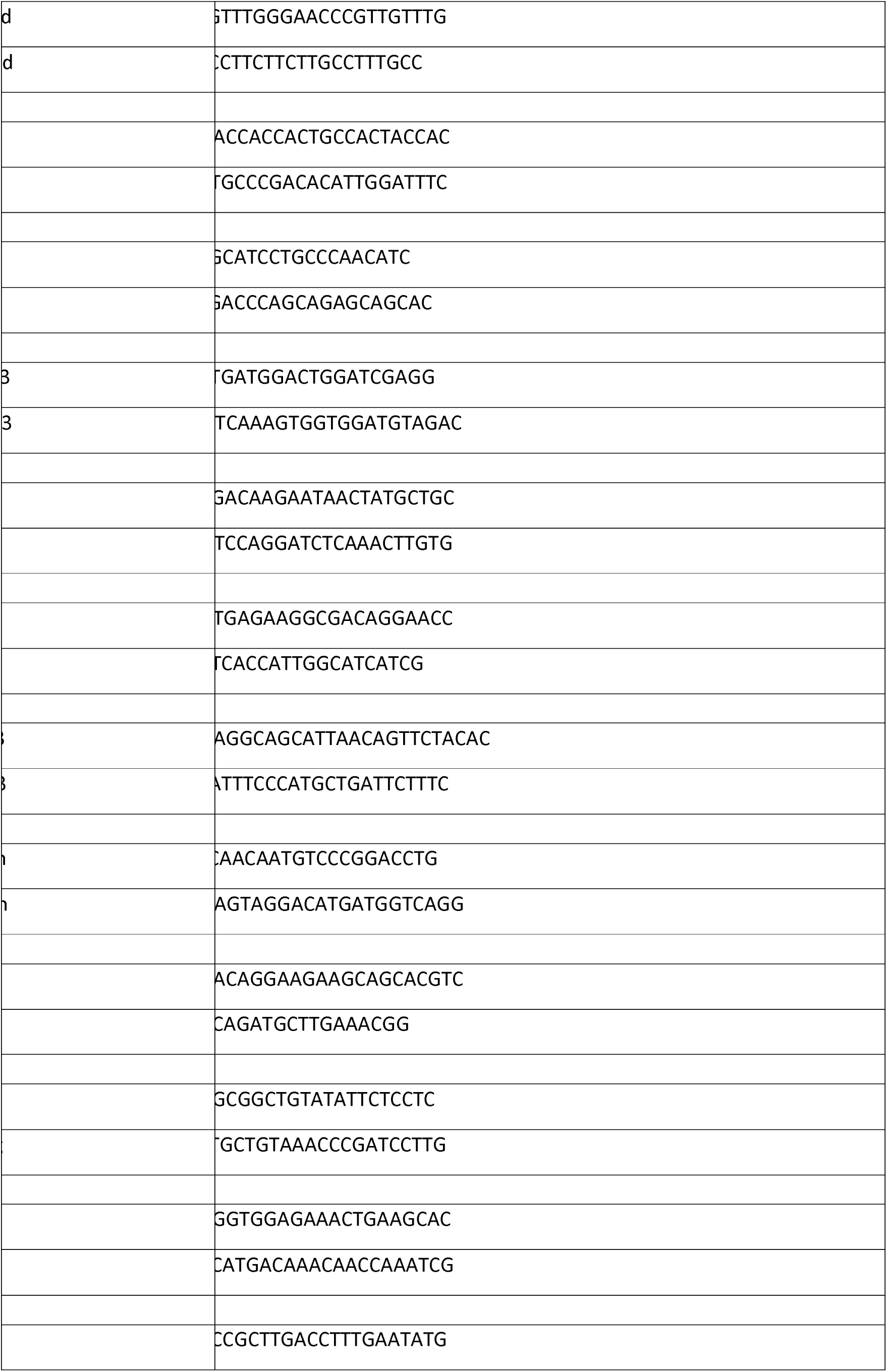
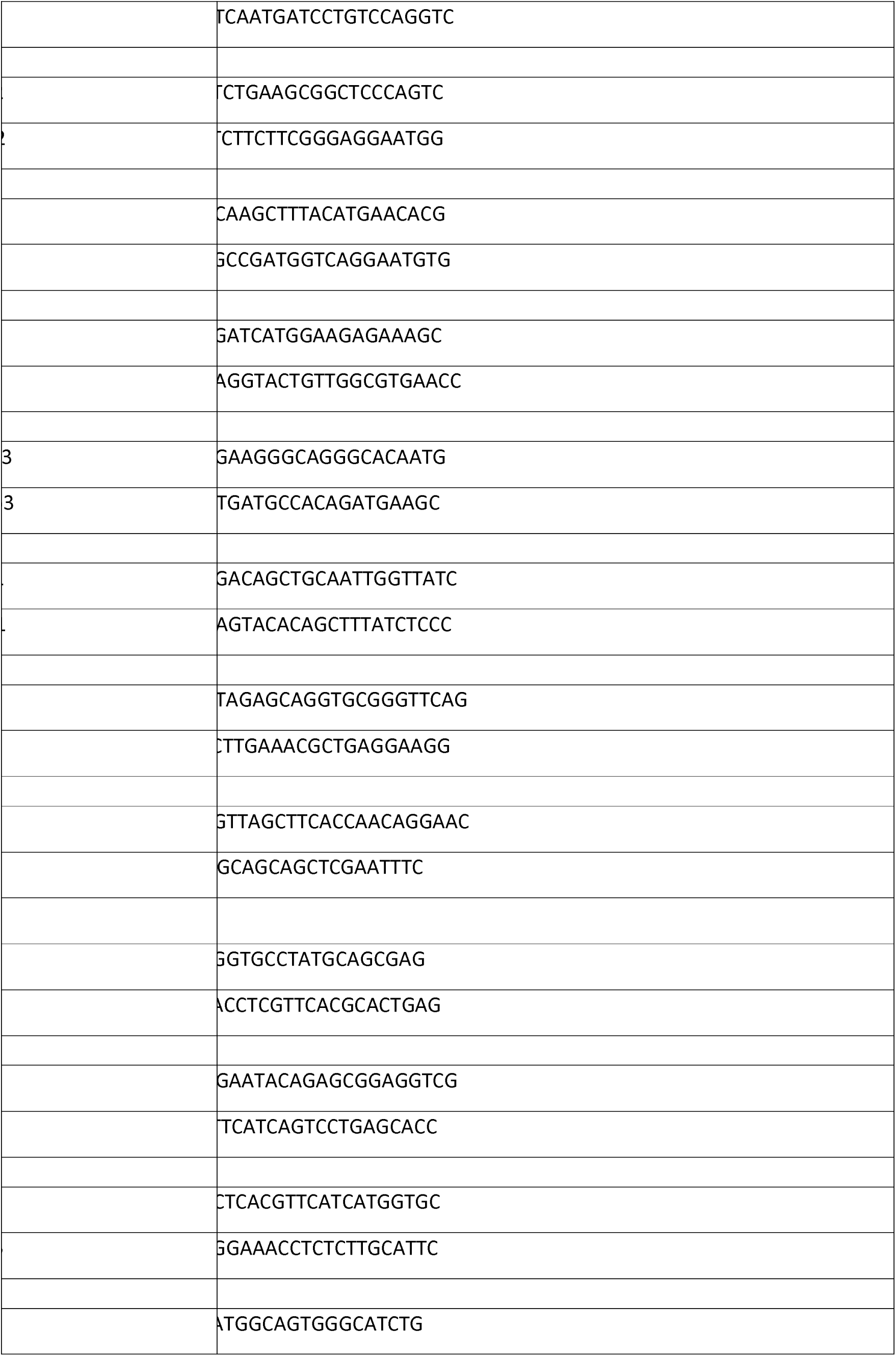
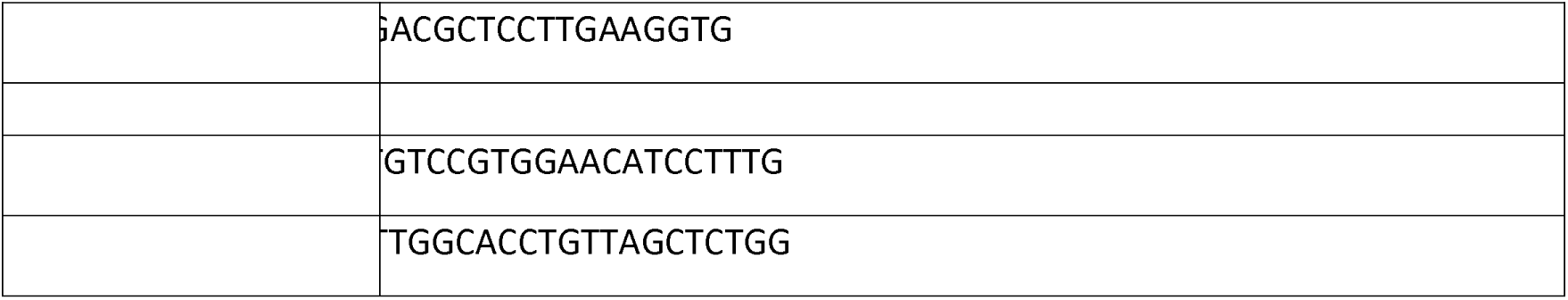
Primers sequences.

## Notes

### Competing Interest Statement

The authors have declared no competing interest.

